# Guiding treatment response by spatiotemporal control of α-particle deposition in solid tumors: the case for ‘affinity cocktails’ of antibody-radioconjugates

**DOI:** 10.64898/2025.12.22.696075

**Authors:** Rajiv Nair, Aira Sarkar, Pooja Hariharan, Mihail Kavousanakis, Rohit Chaudhari, Remco Bastiaannet, Kathlyn Gabrielson, Stavroula Sofou

**Author notes:** corresponding author Stavroula Sofou, ChemBE, Johns Hopkins University 3400 North Charles Street, Maryland Hall 116, Baltimore, MD 21218. equally contributing authors (Rajiv Nair, Aira Sarkar, Pooja Hariharan).

## Abstract

Antibody-radioconjugates are leading the investigational targeted alpha-particle (α-particle) therapies for the treatment of solid tumors that do not respond to approved therapies. Yet, there is still treatment failure in the clinic largely attributed to the heterogenous patterns of tumor irradiation by α-particles. Although α-particles are essentially impervious to resistance, attributed to the complex double-strand DNA breaks they cause while traversing cells, cells not being directly hit by α-particles will likely not be killed. The diffusion-limited poor tumor penetration of high-affinity (strongly-binding) antibody-radioconjugates combined with α-particles’ short-range in tissue (only 40-80μm), let tumor regions far from vasculature inadequately irradiated, therefore, possibly escaping treatment.

**METHODS:** To improve penetration of delivered activity within tumors, we engineered separate actinium-225 antibody-radioconjugates of variable affinities (‘affinity cocktails’) targeting the same marker on cancer cells, that were chosen based on their preferential irradiation of complementary regions of the same tumors. The cocktails comprise: (a) ‘high-affinity’ antibody-radioconjugates (as the ones on clinical trials), which mostly deliver their cargo in tumor cells close to the vasculature, where the ‘low(er)-affinity’ antibody-radioconjugates fail to deliver effective doses, due to their fast clearance; and (b) ‘low(er)-affinity’ antibody-radioconjugates, that penetrate the deeper parts of tumors farther from the vasculature, where the ‘high-affinity’ antibodies fail to reach. The efficacy of affinity cocktails was assessed in spheroids, that were employed as surrogates of tumor avascular regions, and on mice with subcutaneous xenografts of different cancer origin, expression levels and/or type of the targeted receptor: HER2 highly-expressing BT-474 breast cancer cells, HER2 moderately-expressing HEPG2 hepatoma cells, and/or HER1 low-expressing BxPC-3 pancreatic cancer cells.

**RESULTS:** Although the high-affinity antibody-radioconjugates were most lethal against cancer cells in monolayers, affinity cocktails were most effective in inhibiting spheroid growth, due to better collective spreading of the antibody-conjugates within the spheroids’ volume. On all mouse models, and for the same total injected activity, affinity cocktails resulted in the best tumor growth inhibition, even at lower tumor absorbed doses, compared to the high-affinity antibody-radioconjugates alone.

**CONCLUSIONS:** This proof-of-concept study in α-particle antibody-delivery to solid tumors demonstrates that ‘separating’ the two key processes of diffusion and reaction/binding improves treatment efficacy. This generalizable approach may augment antibody-radioconjugates already in clinical trials.

## Introduction

Antibody-radioconjugates are among the leading approaches in targeted <u>a</u>lpha-particle (α-particle) radionuclide therapies (TAT) that are currently being clinically evaluated against solid tumors of different origins (*1*). The enthusiasm for TAT is due to the unparalleled killing efficacy of, and irradiation precision by, α-particles and the selectivity in tumor targeting by antibody technologies (*1*). α-particles travel only 4-5 cell lengths in tissue, and, due to their high energy, they cause complex double-strand DNA breaks (*2*). The inability to repair this DNA damage is the reason that α-particles (*2*) are largely impervious to resistance (*3–5*), *independent of cell origin and/or resistance to other agents* (*3*). However, cells not being directly hit by α-particles will likely not be killed. Antibody-radioconjugates targeting markers on cancer cells are successful in selectively delivering significant doses to tumors, but the more strongly an antibody binds on cancer cells, the less deep it penetrates into solid tumors (*6,7*); the limited penetration in solid tumors of high-affinity antibody-radioconjugates results in fewer cancer cells being directly irradiated and killed. Limited penetration leading to heterogeneous tumor irradiation is a main reason why patients treated with TAT often ultimately relapse (*8*).

In this study, we aimed to improve the efficacy of α-particle antibody-radioconjugates, by enabling deep penetration of α-particles in solid tumors. Our approach employs antibody-radioconjugates of *controlled affinities (“affinity cocktails”).* We simultaneously deliver the same α-particle emitter by separate antibodies, each preferentially irradiating different regions of the tumor: (1) a high-affinity antibody-radioconjugate (identical to the ones currently in clinical trials) irradiating mostly the tumor perivascular regions (where aggressively growing cancer cells reside); and (2) a separately administered low(er)-affinity antibody-radioconjugate that upon tumor uptake penetrates the deeper parts of tumors where high-affinity antibodies *do not reach* (and where cancer recurrence originates from). Although the low(er)-affinity antibodies clear rapidly from the tumor perivascular regions, since they do not strongly bind to cells (*9*), these perivascular regions are effectively being irradiated and killed by the high-affinity radiolabeled-antibodies.

As a proof-of-concept, on mice with subcutaneous xenografts of liver cancer with moderate expression of the human epidermal growth factor receptor 2 (HER2), we assessed the efficacy to inhibit tumor growth of an affinity cocktail comprised an actinium-225 HER2-targeting trastuzumab-radioconjugate in its regular “high-affinity” form (high-affinity Ab-radioconjugate) and in a low(er)-affinity form (low-affinity Ab-radioconjugate) and compared to each antibody-radioconjugate alone, at same total injected activity.

To demonstrate the general applicability of affinity cocktails in improving tumor growth inhibition relative to high-affinity antibody-radioconjugates alone, we assessed this approach on mice with subcutaneous xenografts of different origin (breast, liver, and pancreatic cancer) expressing high, moderate and low levels of the targeted receptor (HER2 or HER1). We employed trastuzumab-radioconjugates or cetuximab-radioconjugates as the corresponding high-affinity antibody-radioconjugates, and we chose rituximab (a CD20-targeting antibody, which was not expected to recognize any markers on cancer cells) employed as a *model low-affinity antibody-radioconjugate that was common across all tumor types*.

## Materials and Methods

Indium-111 (^111^In) chloride was purchased from BWXT (Ontario, Canada). The Actinium-225 used in this research was supplied by the U.S. Department of Energy Isotope Program, managed by the Office of Isotope R&D and Production. Information on other reagents is detailed in the supporting information section.

### Cell lines

The cell lines BT-474 and HEPG2 were purchased from ATCC and were cultured using Hybricare^TM^ Media and Eagle’s Minimum Essential Medium (EMEM), respectively, each supplemented with 10% FBS, 100 units/mL Penicillin and 100µg/mL Streptomycin at 37°C and 5% CO_2_. The cell line BxPC-3 was a gift from Dr Denis Wirtz at JHU and was cultured in Roswell Park Memorial Institute (RPMI) media, supplemented as above.

### Lowering Trastuzumab’s affinity

Fluorescein (FITC)-NHS (5 mg/mL in DMSO) was added dropwise on ice to trastuzumab (in PBS, pH 7.4), at a molar ratio of 50:1 (fluorescein-to-antibody) and was incubated overnight to selectively react with the α-amino groups at the N-terminus (paratopes). Low-affinity trastuzumab-FITC was purified by a 10DG column equilibrated with 0.1M sodium carbonate buffer pH 9.0, for further conjugation with DOTA-SCN, and its immunoreactivity was measured on BT-474 cells in excess of HER2-receptors, as previously reported (*10*).

### Antibody chelator/fluorophore conjugation in the Fc region

DOTA-SCN (DTPA-SCN or FITC-SCN; dissolved in DMF or DMSO at 10-20mg/mL) was added on ice to antibodies (2.5mg/mL in 0.1M sodium carbonate buffer, pH 9.0), and the reaction was allowed to proceed overnight followed by a 10DG column that was used for purification.

For radiolabeling, activity (in 0.2 N HCl) was added to the corresponding antibody solutions following 1 hr incubation at 37°C at final pH=9 (*6*). A 10DG column was employed for the antibody-radioconjugate purification. Radiolabeling efficiency, radiochemical purity, the immunoreactivity of the antibody-radioconjugate, and the stability of radiolabeling were evaluated as previously reported (*6,11*) and detailed in the supporting information section. The antibody concentration was measured using the BCA assay.

### Clonogenic cell survival assay

Cells in monolayers were exposed for 6 hours to actinium-225 (delivered by different carriers, as indicated) and were then plated in tissue culture dishes till the onset of colony formation (∼10 cell doubling times). The clonogenic cell survival fractions were counted as previously described (*12*) and detailed in the supporting information section.

### Flow cytometry

Cells were trypsinized, were suspended in media (at 1 million cells/mL), and were incubated on ice for 1 hour with FITC-labeled antibody-conjugates (at 50 x 10^6^ antibody-to-cell ratio). The cell suspensions were then centrifuged thrice, were resuspended in ice cold PBS and were analyzed at the same settings (450 mV) on the BD FACS Canto Flow cytometer (Franklin Lakes, NJ, USA).

### Spheroids

Spheroids were formed by seeding 3,200 BT-474 or 1,000 HEPG2 cells per well in a PolyHEMA-coated 96-well round bottom plate, followed by centrifugation for 10 minutes at 1,023rcf and 4°C. The formed spheroids were tracked for their size and were used upon reaching 400µm-in-diameter. The cell line BxPC-3 did not form spheroids.

To evaluate the antibodies’ spatiotemporal distributions in spheroids, spheroids were incubated with FITC-labeled antibody-conjugates (0.06µM) in150μL and at various timepoints, spheroids were sampled, flash frozen in cryochrome, mounted on OCT gel and sectioned at 20µm thickness. The equatorial sections were then imaged (ex/em: 494/518nm) using a confocal fluorescence microscope (Zeiss LSM 780 Confocal Microscope with filters, 10X objective, White Plains, NY). The calibration curves of the fluorescent antibody-conjugates were generated using a 20µm-pathlength quartz cuvette measured on the same instrument. The mean radial concentrations of antibody-conjugates in spheroid sections were calculated using an in-house erosion code averaging 5 µm-wide concentric rings.

In treatment studies, spheroids were incubated for 24 hours with varying combinations of antibody-radioconjugates at different activity concentrations. The total antibody mass was maintained at 10 µg/mL in all cases, corresponding to approximately 300 times excess of the HER2 receptors expressed by all cells comprising the spheroid. After incubation, spheroids were transferred into fresh media (one spheroid per well in PolyHEMA-coated U-bottom plates) and were allowed to grow for 10 more days before being transferred to adherent 96-well plates (one spheroid per well). Once the non-treated condition reached ∼80% confluency, cells were trypsinized and counted. The percentage regrowth was evaluated as the number of cells counted in each treated condition normalized by the number of cells in the untreated condition.

### Animal study

All animal studies were performed in compliance with Institutional Animal Care and Use Committee protocol (IACUC) guidelines. Four-to-six weeks old, 20g NSG (NOD SCID-gamma) mice were purchased from JHU Breeding Facility. Mice were housed in filter-top cages with sterile food and water.

Tumor xenografts were established subcutaneously by injecting inoculums of 100µL (50:50 v/v ratio of serum-free medium and Matrigel^TM^) containing 1,000,000 BT-474 cells per female mouse, 1,500,000 HEPG2 cells or 500,000 BxPC-3 cells per male mouse. When tumor volumes reached approximately 100mm^3^, animals were randomly assigned to a group, and each mouse was administered intravenously a total of 2.96kBq actinium-225 delivered by antibody-radioconjugates as indicated. Tumor size was monitored and measured by a digital caliper (resolution: 0.01 mm), and animal weights were recorded. The tumor volumes were calculated using V=4*π*α*β^2^/3, where α and β are the major and minor diameters, respectively. The endpoint criterion was set as the point when tumor volume reached 400mm^3^. Following euthanasia, excised tumors and critical organs were fixed and H&E stained for histological evaluation.

### Dosimetry

Dosimetry was calculated using the software package 3D-RD-S, Radiopharmaceutical Imaging and Dosimetry, LLC (Rapid, Baltimore, MD) from the biodistributions of intravenously injected [^111^In]In-antibody-radioconjugates (740kBq per mouse) on the corresponding tumor-bearing mouse models, as previously reported (*6,11*). In dosimetry calculations, all α-particles’ and electron energies delivered by the antibodies were assumed to be absorbed by the tissue where the first decay occurred (*6,11*).

### Statistical Analysis

Data were reported as the arithmetic mean of n independent measurements ± the standard deviation. To calculate the differences between experimental groups, one-way ANOVA and/or the unpaired Student’s t test were employed. Kaplan-Meier survival plots were analyzed using the log-rank test. p-values<0.05 were considered significant. * indicates 0.01<*p*-values<0.05; **<0.01, ***<0.001.

## Results

### Antibody and cell line characterization

Table 1 demonstrates stable and reproducible radiolabeling both with [^225^Ac]Ac and with [^111^In]In for all antibodies studied: (high-affinity) trastuzumab radioconjugates, low-affinity trastuzumab radioconjugates, (high-affinity) cetuximab radioconjugates, and the common *model* low/no-affinity antibody-radioconjugate (rituximab-radioconjugate).

**Table 1.**
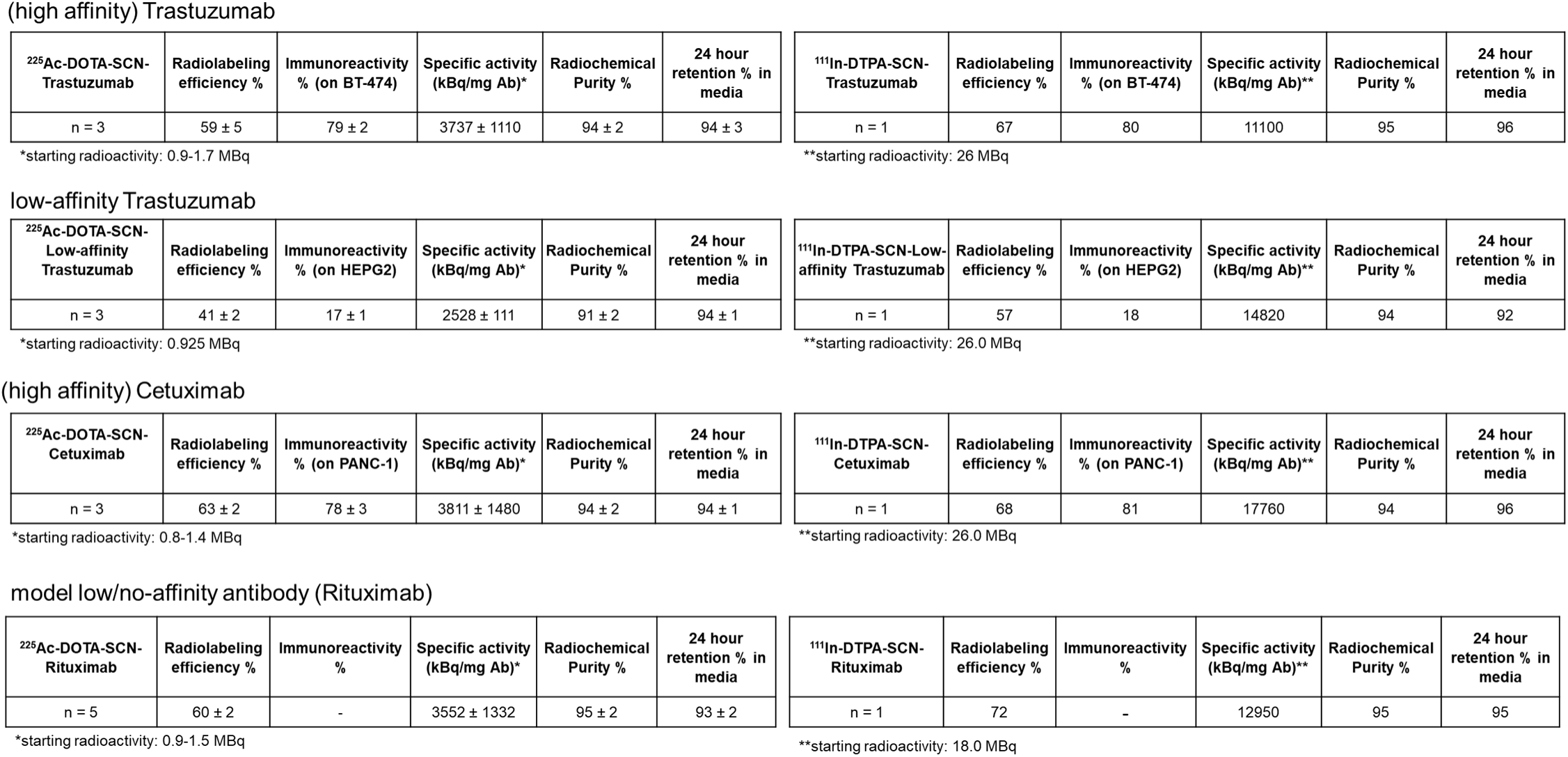
Characterization of antibody-radioconjugates. Shown are the mean values ± the standard deviation of n independent measurements. Stability was assessed as % retention of activity by antibody-radioconjugates following a 24-hour incubation in complete media at 37°C.

(high affinity) Trastuzumab
| $^{225}\text{Ac}$-DOTA-SCN-Trastuzumab | Radiolabeling efficiency % | Immunoreactivity % (on BT-474) | Specific activity (kBq/mg Ab)* | Radiochemical Purity % | 24 hour retention % in media | $^{111}\text{In}$-DTPA-SCN-Trastuzumab | Radiolabeling efficiency % | Immunoreactivity % (on BT-474) | Specific activity (kBq/mg Ab)** | Radiochemical Purity % | 24 hour retention % in media |
| --- | --- | --- | --- | --- | --- | --- | --- | --- | --- | --- | --- |
| n = 3 | 59 ± 5 | 79 ± 2 | 3737 ± 1110 | 94 ± 2 | 94 ± 3 | n = 1 | 67 | 80 | 11100 | 95 | 96 |

\*starting radioactivity: 0.9-1.7 MBq
\*\*starting radioactivity: 26 MBq

low-affinity Trastuzumab
| $^{225}\text{Ac}$-DOTA-SCN-Low-affinity Trastuzumab | Radiolabeling efficiency % | Immunoreactivity % (on HEPG2) | Specific activity (kBq/mg Ab)* | Radiochemical Purity % | 24 hour retention % in media | $^{111}\text{In}$-DTPA-SCN-Low-affinity Trastuzumab | Radiolabeling efficiency % | Immunoreactivity % (on HEPG2) | Specific activity (kBq/mg Ab)** | Radiochemical Purity % | 24 hour retention % in media |
| --- | --- | --- | --- | --- | --- | --- | --- | --- | --- | --- | --- |
| n = 3 | 41 ± 2 | 17 ± 1 | 2528 ± 111 | 91 ± 2 | 94 ± 1 | n = 1 | 57 | 18 | 14820 | 94 | 92 |

\*starting radioactivity: 0.925 MBq
\*\*starting radioactivity: 26.0 MBq

(high affinity) Cetuximab
| $^{225}\text{Ac}$-DOTA-SCN-Cetuximab | Radiolabeling efficiency % | Immunoreactivity % (on PANC-1) | Specific activity (kBq/mg Ab)* | Radiochemical Purity % | 24 hour retention % in media | $^{111}\text{In}$-DTPA-SCN-Cetuximab | Radiolabeling efficiency % | Immunoreactivity % (on PANC-1) | Specific activity (kBq/mg Ab)** | Radiochemical Purity % | 24 hour retention % in media |
| --- | --- | --- | --- | --- | --- | --- | --- | --- | --- | --- | --- |
| n = 3 | 63 ± 2 | 78 ± 3 | 3811 ± 1480 | 94 ± 2 | 94 ± 1 | n = 1 | 68 | 81 | 17760 | 94 | 96 |

\*starting radioactivity: 0.8-1.4 MBq
\*\*starting radioactivity: 26.0 MBq

model low/no-affinity antibody (Rituximab)
| $^{225}\text{Ac}$-DOTA-SCN-Rituximab | Radiolabeling efficiency % | Immunoreactivity % | Specific activity (kBq/mg Ab)* | Radiochemical Purity % | 24 hour retention % in media | $^{111}\text{In}$-DTPA-SCN-Rituximab | Radiolabeling efficiency % | Immunoreactivity % | Specific activity (kBq/mg Ab)** | Radiochemical Purity % | 24 hour retention % in media |
| --- | --- | --- | --- | --- | --- | --- | --- | --- | --- | --- | --- |
| n = 5 | 60 ± 2 | - | 3552 ± 1332 | 95 ± 2 | 93 ± 2 | n = 1 | 72 | - | 12950 | 95 | 95 |

\*starting radioactivity: 0.9-1.5 MBq
\*\*starting radioactivity: 18.0 MBq

The mean expression levels of targeted receptors per cell line, measured by (high-affinity) trastuzumab-FITC were 1,200,000 ± 24,000 HER2-copies per BT-474 and 410,000 ± 12,000 HER2-copies per HEPG2, and 117,000 ± 6,000 HER1-copies per BxPC-3 measured by (high-affinity) cetuximab-FITC (as shown by the binding isotherms in Figure S1). The dissociation constant (K_D_) of (high-affinity) trastuzumab ranged from 1.6 to 7.0 nM, and of (high-affinity) cetuximab was 0.3 nM.

In all Figures, red indicates the corresponding high-affinity (forms of) antibodies, blue the low(er) affinity antibodies, and purple or pink the affinity cocktails with activity split ratios equal to 50:50 and 70:30 between the high- and low-affinity radioconjugates, respectively.

On all three cell lines studied, flow cytometry confirmed the greater fluorescence shift by each of the two high-affinity antibodies (shown in red: the HER2-targeting trastuzumab in Figures 1A, 1B, 1C; and the HER1-targeting cetuximab in Figure 1D), compared to the minor shifts of the corresponding low-affinity antibodies (shown in blue: the HER2-targeting low-affinity trastuzumab in Figure 1A; and the model low/no-affinity rituximab in Figures 1B, 1C, 1D).

**FIGURE 1.**
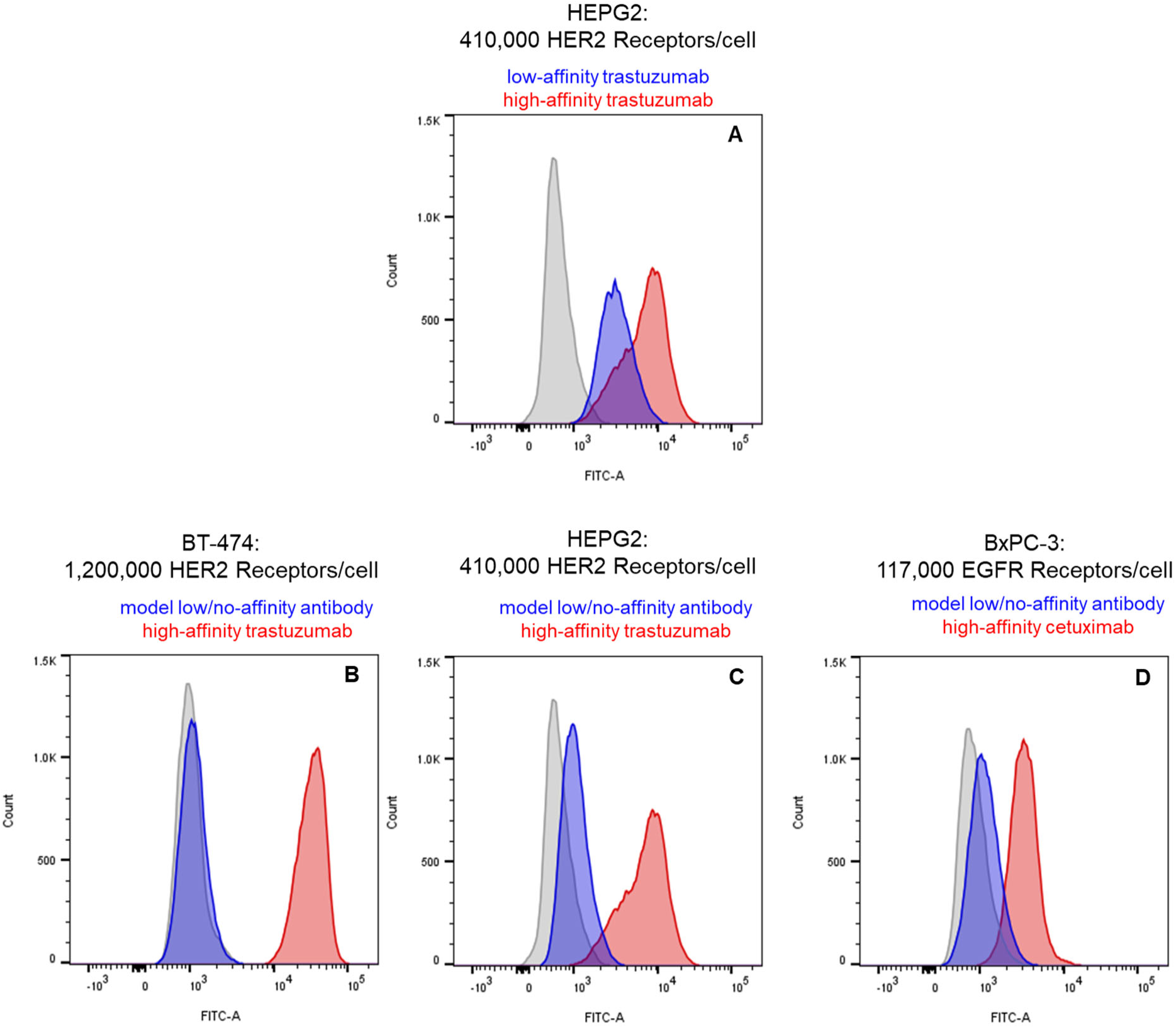
Flow cytometry of different cell types incubated with the corresponding FITC-labeled antibodies of different affinities. Red: (high-affinity) trastuzumab, (high affinity) cetuximab; Blue: low-affinity trastuzumab or the common *model* low/no-affinity antibody (rituximab). Gray: cells only.

### In vivo studies: dosimetry

Dosimetry for all [²²⁵Ac]Ac-DOTA-antibodies on the three mouse models (Table 2) was calculated using the biodistributions of the corresponding, systemically administered [¹¹¹In]In-DTPA-labeled antibodies (see Tables S1-S3 and Figures S2-S4).

**Table 2.**
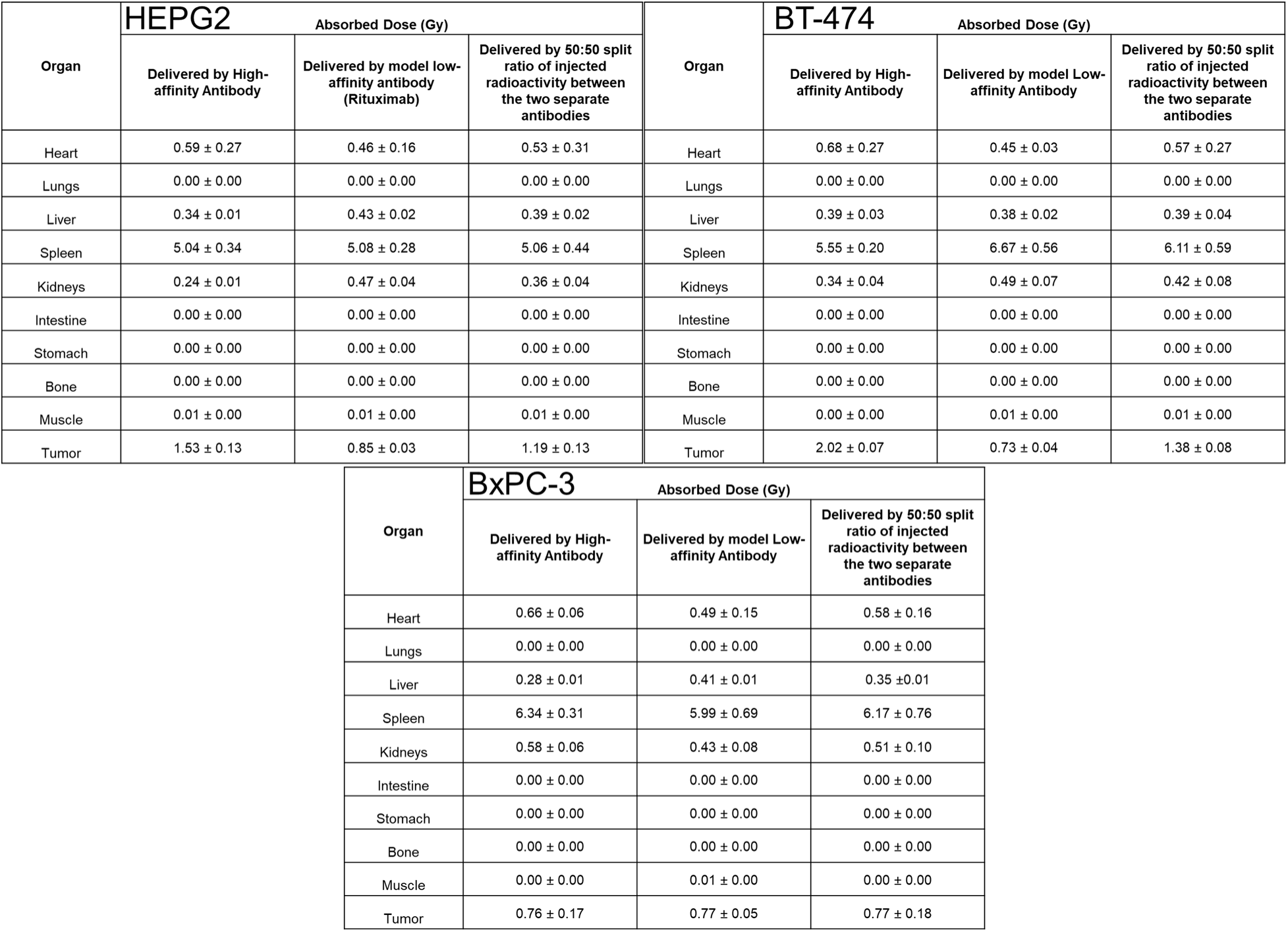
Dosimetry calculations were based on the administered activities of 2.96kBq [²²⁵Ac]Ac-DOTA-antibody conjugates per mouse. Mean absorbed doses ± standard deviations are reported.

| HEPG2 |  |  |  | Organ | BT-474 |  |  |
| --- | --- | --- | --- | --- | --- | --- | --- |
| Organ | Delivered by High-affinity Antibody | Delivered by model low-affinity antibody (Rituximab) | Delivered by 50:50 split ratio of injected radioactivity between the two separate antibodies |  | Delivered by High-affinity Antibody | Delivered by model Low-affinity Antibody | Delivered by 50:50 split ratio of injected radioactivity between the two separate antibodies |
| Heart | 0.59 ± 0.27 | 0.46 ± 0.16 | 0.53 ± 0.31 | Heart | 0.68 ± 0.27 | 0.45 ± 0.03 | 0.57 ± 0.27 |
| Lungs | 0.00 ± 0.00 | 0.00 ± 0.00 | 0.00 ± 0.00 | Lungs | 0.00 ± 0.00 | 0.00 ± 0.00 | 0.00 ± 0.00 |
| Liver | 0.34 ± 0.01 | 0.43 ± 0.02 | 0.39 ± 0.02 | Liver | 0.39 ± 0.03 | 0.38 ± 0.02 | 0.39 ± 0.04 |
| Spleen | 5.04 ± 0.34 | 5.08 ± 0.28 | 5.06 ± 0.44 | Spleen | 5.55 ± 0.20 | 6.67 ± 0.56 | 6.11 ± 0.59 |
| Kidneys | 0.24 ± 0.01 | 0.47 ± 0.04 | 0.36 ± 0.04 | Kidneys | 0.34 ± 0.04 | 0.49 ± 0.07 | 0.42 ± 0.08 |
| Intestine | 0.00 ± 0.00 | 0.00 ± 0.00 | 0.00 ± 0.00 | Intestine | 0.00 ± 0.00 | 0.00 ± 0.00 | 0.00 ± 0.00 |
| Stomach | 0.00 ± 0.00 | 0.00 ± 0.00 | 0.00 ± 0.00 | Stomach | 0.00 ± 0.00 | 0.00 ± 0.00 | 0.00 ± 0.00 |
| Bone | 0.00 ± 0.00 | 0.00 ± 0.00 | 0.00 ± 0.00 | Bone | 0.00 ± 0.00 | 0.00 ± 0.00 | 0.00 ± 0.00 |
| Muscle | 0.01 ± 0.00 | 0.01 ± 0.00 | 0.01 ± 0.00 | Muscle | 0.00 ± 0.00 | 0.01 ± 0.00 | 0.01 ± 0.00 |
| Tumor | 1.53 ± 0.13 | 0.85 ± 0.03 | 1.19 ± 0.13 | Tumor | 2.02 ± 0.07 | 0.73 ± 0.04 | 1.38 ± 0.08 |

| Organ | BxPC-3 |  |  |
| --- | --- | --- | --- |
|  | Absorbed Dose (Gy) |  |  |
|  | Delivered by High-affinity Antibody | Delivered by model Low-affinity Antibody | Delivered by 50:50 split ratio of injected radioactivity between the two separate antibodies |
| Heart | 0.66 ± 0.06 | 0.49 ± 0.15 | 0.58 ± 0.16 |
| Lungs | 0.00 ± 0.00 | 0.00 ± 0.00 | 0.00 ± 0.00 |
| Liver | 0.28 ± 0.01 | 0.41 ± 0.01 | 0.35 ±0.01 |
| Spleen | 6.34 ± 0.31 | 5.99 ± 0.69 | 6.17 ± 0.76 |
| Kidneys | 0.58 ± 0.06 | 0.43 ± 0.08 | 0.51 ± 0.10 |
| Intestine | 0.00 ± 0.00 | 0.00 ± 0.00 | 0.00 ± 0.00 |
| Stomach | 0.00 ± 0.00 | 0.00 ± 0.00 | 0.00 ± 0.00 |
| Bone | 0.00 ± 0.00 | 0.00 ± 0.00 | 0.00 ± 0.00 |
| Muscle | 0.00 ± 0.00 | 0.01 ± 0.00 | 0.00 ± 0.00 |
| Tumor | 0.76 ± 0.17 | 0.77 ± 0.05 | 0.77 ± 0.18 |

### Cocktails of high-affinity and low-affinity HER2-targeting trastuzumab-radioconjugates

For moderately HER-expressing HEPG2 hepatoma cells in monolayers, in the absence of diffusion-limited access to cells, Fig. 2A shows that actinium-225 was most lethal when delivered by the (high-affinity) trastuzumab radioconjugate (red symbols) followed by the affinity cocktail (purple symbols, indicating equal activity split between the two trastuzumab forms). However, Fig. 2B shows that in 3D spheroids, that capture the diffusion-limited antibody-penetration (*6,10*), the lowest extents of spheroid regrowth were observed after treatment with affinity cocktails (for activity split ratios ranging from 30:70 to 70:30 between the two affinity-forms of antibody-radioconjugates), at the same total actinium-225 activity concentration during incubation. This outcome was attributed to the complementary microdistributions within the spheroids of each of the two affinity-forms of antibodies. In particular, Fig. 2C shows that (a) close to the spheroid edge, the (high-affinity) trastuzumab (red symbols, red arrow) exhibited higher localization relative to the localization of the low-affinity trastuzumab (blue symbols, blue arrow); and (b) in the spheroid core, the low-affinity trastuzumab (blue symbols, blue bracket) penetrated at a greater extent than the high-affinity trastuzumab (red symbols, red bracket). The spatiotemporal profiles of the two affinity-forms of the trastuzumab-conjugates are shown in Figure S5.

**Figure 2.**
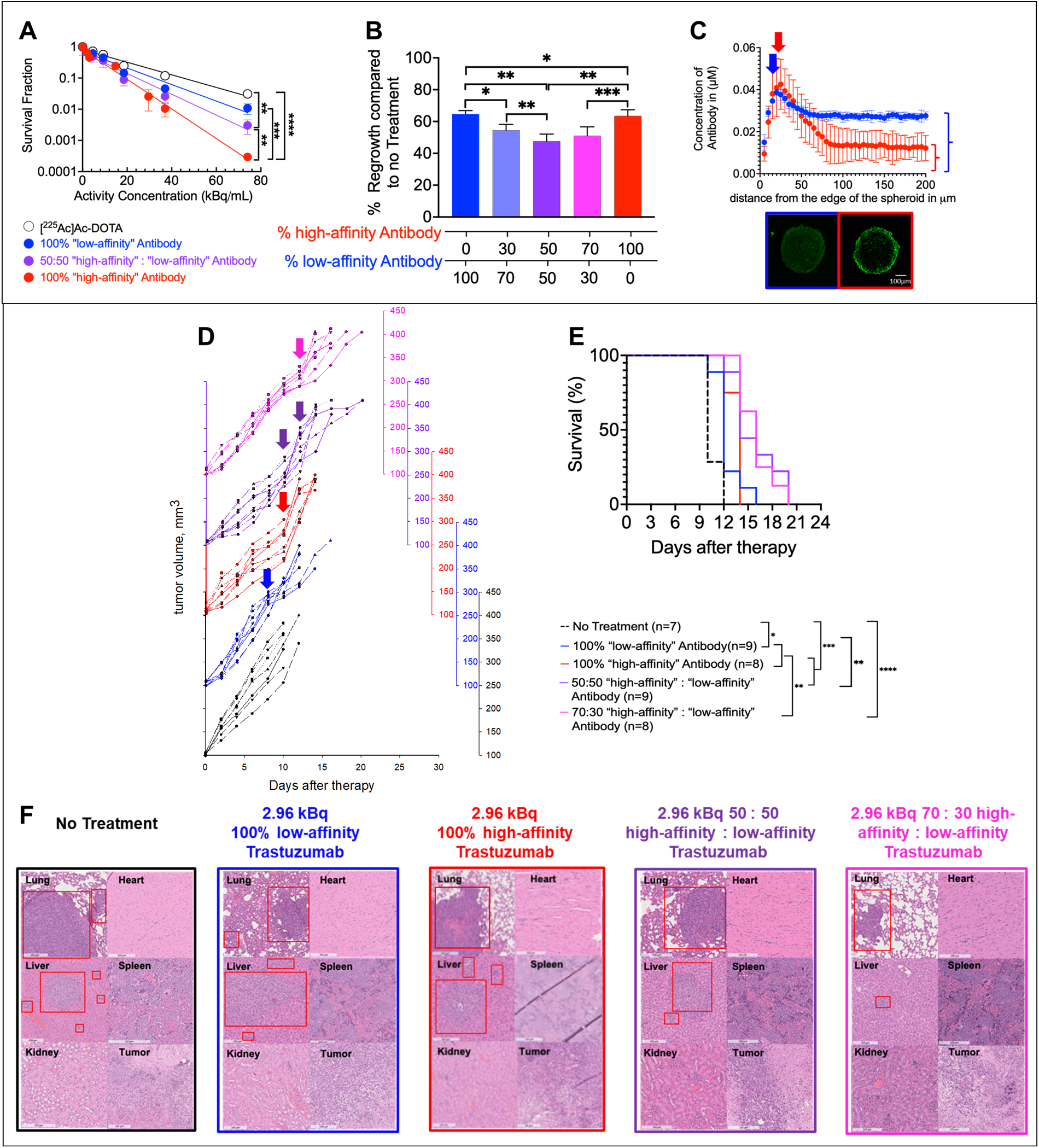
HEPG2 hepatoma cells with moderate HER2-expression: Cocktails of high and low affinity HER2-targeting trastuzumab-radioconjugates best inhibit tumor growth and result in the lowest hepatic metastatic burden. (A) Clonogenic survival fraction following exposure to actinium-225 delivered by a (high-affinity) trastuzumab-radioconjugate (red symbols), a low-affinity trastuzumab-radioconjugate (blue symbols), and/or by equally splitting the activity between both antibody-radioconjugates (purple symbols). (B) Extent of HEPG2 spheroid regrowth, compared to non-treated, after exposure to 1 kB/mL actinium-225 split into different ratios between the high/low-affinity trastuzumab-radioconjugates. (C) Fluorescent microscopy images and radial antibody concentrations in equatorial sections of spheroids, employed as surrogates of tumors’ avascular regions, following after 24 hours of incubation. (D) Growth inhibition of subcutaneous HEPG2 xenografts in NSG mice injected systemically with 2.96kBq actinium-225 delivered by each affinity form of trastuzumab-radioconjugate alone and as affinity cocktails at two different activity split ratios. Vertical arrows indicate the inflection points beyond which tumors exhibited increased growth rates. (E) Animal survival indicating the time required for tumor volumes to reach/not exceed the end-point of 400mm^3^. (F) H&E staining of tumors and critical off-target organs: The 70:30 affinity cocktail (pink frame) demonstrated the lowest hepatic metastatic burden (indicated by red boxes) among all groups. Scale bar = 200µm.

On mice (Fig. 2D-F), almost all systemically treated HEPG2 xenografts (Fig. 2D) exhibited two growth phases: an initial (slower) rate phase followed by an inflection point (indicated by the vertical arrows) beyond which the growth rate generally increased. All tumor-bearing mice were treated with the same total injected activity of 2.96kBq, that was previously shown to not cause off-target toxicities (*3*). Even at lower tumor absorbed doses (Figure S2), compared to the high-affinity antibody-radioconjugate alone (red symbols), the affinity-cocktail at 70:30 activity-split ratio (pink symbols) not only delayed the inflection point by two days (day 12 vs day 10 after initiation of treatment), but, importantly, resulted in the lowest median tumor growth rate after the inflection point (19 mm^3^/day vs 36 mm^3^/day, respectively). The affinity-cocktail of 50:50 activity-split ratio (purple symbols) exhibited a bimodal behavior with a fraction of the tumors behaving similarly to tumors after the 70:30 affinity-cocktail treatment (pink symbols) and the rest similarly to the response after the high-affinity antibody-radioconjugate treatment (red symbols). Animal survival in Fig. 2E indicates the time required for tumor volumes to reach/not exceed the end-point of 400mm^3^.

Pathology evaluation, shown in Fig. 2F, across all cohorts revealed no evidence of treatment-related cardiac, renal, or hepatic toxicity. Importantly, the 70:30 affinity cocktail (pink frame) produced extensive necrosis, along with mineral deposition consistent with acidic tumor microenvironments, and demonstrated the lowest hepatic metastatic burden among the groups. The 50:50 affinity cocktail (purple frame) induced a more pronounced increase in necrosis compared to non-treated, accompanied by EMH in liver and spleen, without evidence of organ toxicity. The (high-affinity) antibody-radioconjugate cohort (red frame) exhibited fewer lung lesions and a greater extent of necrotic regions relative to non-treated. The low-affinity antibody-radioconjugate cohort (blue frame) showed invasive tumor growth into skeletal muscle with moderate necrosis, with fewer lung and liver metastases than non-treated, but less cell killing efficacy than the affinity-cocktail antibody-radioconjugates. Tumors were uniformly large, with spindle-shaped cells, and showed areas of necrosis. Spleens in all groups displayed prominent extramedullary hematopoiesis (EMH). In tumors receiving α-particle therapy, an EMT-like phenotype was retained; however, these tumors exhibited greater areas of cell death relative to non-treated and reduced metastatic burden in the liver and lungs (indicated by the red boxes). (animal weight measurements vs. time are shown in Fig. S6).

### Affinity cocktails of antibody-radioconjugates as a general approach

On mice with HER2-overexpressing BT-474, HER2-moderately expressing HEPG2 or low HER1-expressing BxPC-3 xenografts, a single common *model* low-affinity antibody-radioconjugate (rituximab-radioconjugate) was employed for all tumor models, in this proof-of-concept study, and, a single affinity cocktail at 50:50 activity split ratio between the antibody forms was chosen. Fig. 3A shows the same killing efficacy trends on cell monolayers, as in Fig. 2A, with the high-affinity antibody-radioconjugates (HER2-targeting and HER1-targeting, both in red) to result in the least clonogenic survival fractions across all cell lines. On spheroids (shown in Fig. 3B), and in agreement with Fig. 2B, the least spheroid regrowth was exhibited after treatment with the corresponding affinity-cocktails, both in highly and in moderately HER2-expressing spheroids of BT-474 and HEPG2, respectively. These findings correlated with the corresponding microdistributions of each antibody form shown in Fig. 3C, collectively exhibiting complementary patterns within the spheroids as discussed for Fig. 2C. The spatiotemporal profiles of all antibody affinity-forms are shown in Figures S5 and S7.

**Figure 3.**
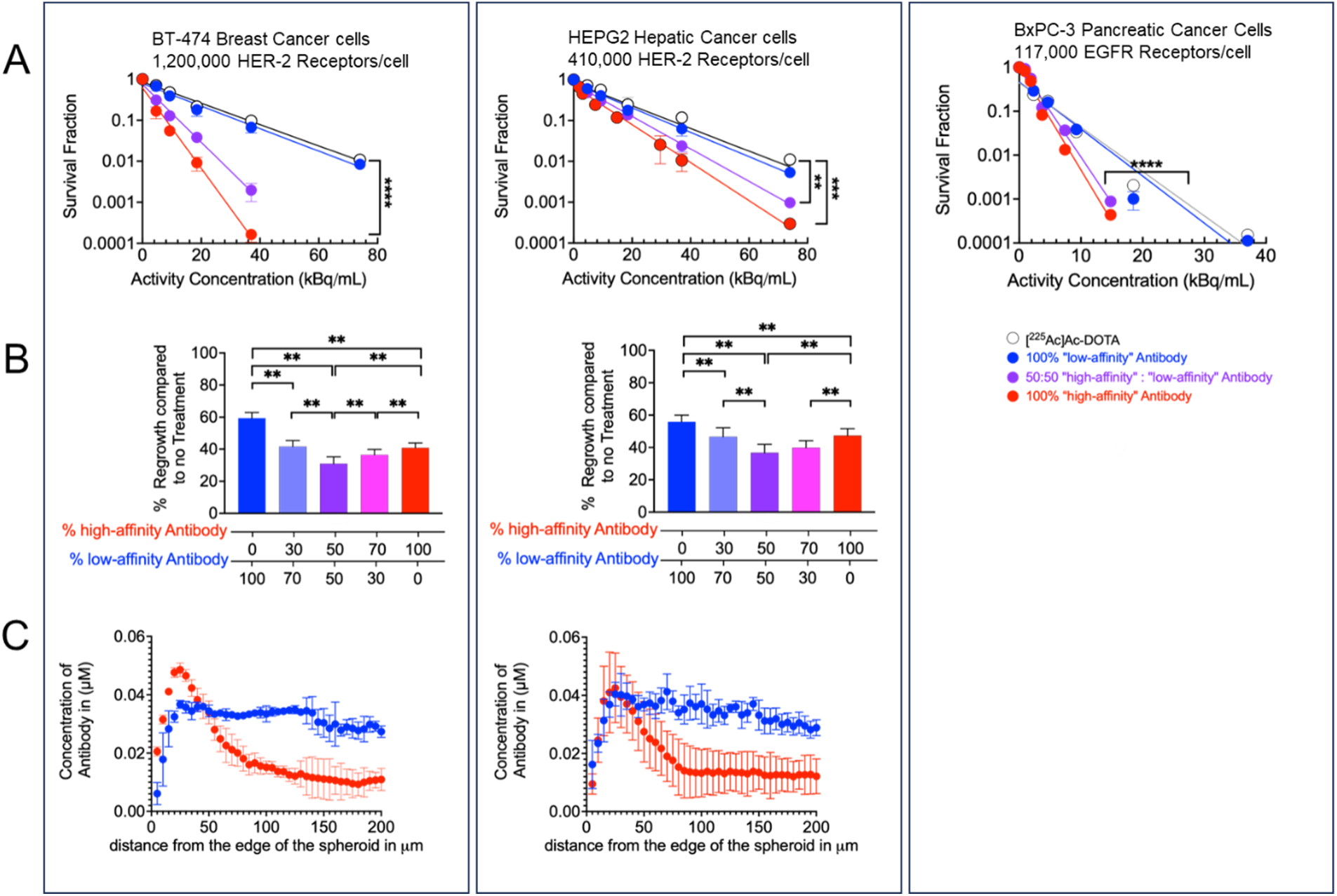
Clonogenic survival fraction (A) and spheroid regrowth (B) after exposure to [^225^Ac]Ac delivered by high-affinity antibody-radioconjugates (red symbols), low-affinity antibody-radioconjugates (blue symbols) and/or their combination. (C) shows the radially averaged concentrations of each antibody type in spheroids following incubation for 24 hours (Fig S5 and S7 show the spatial profiles over time). BxPC-3 cells did not form spheroids. Mean values ± the standard deviation of n=2 (in A), n=3 (in B) and n=3 (in C) independent runs are shown. In all studies the total concentrations of antibodies were kept constant at 10µg/mL (by compensating with addition of the corresponding cold antibodies). Trastuzumab and cetixumab radioconjugates were employed as the high-affinity antibody-radioconjugates for the HER2-expressing BT-474 and HEPG2, and for the HER1-expressing BxPC-3, respectively. Rituximab-radioconjugate was used as a model low/no-affinity antibody-radioconjugate that was common across all tumor lines.

Fig. 4A shows that in all xenografts studied, the same total injected activity when delivered by the affinity cocktails (purple lines) relative to being delivered by the high-affinity antibody alone (red lines): (a) shifted the inflection points toward later days (11 vs. 8, 12 vs. 8-9, and 15 vs. 13 in BT-474, HEPG2, and BxPC-3 xenografts, respectively, as indicated by the purple and red vertical arrows); and (b) decreased the mean tumor growth rates after the inflection point (19 vs. 20, 19 vs. 21, and 15 vs. 19 mm^3^/day in BT-474, HEPG2, and BxPC-3 xenografts, respectively); even (c) at lower tumor absorbed doses (1.2 vs. 1.5 Gy, and 1.4 vs. 2.0 Gy in the BT-474 and HEPG2 xenograft models, respectively) or same tumor absorbed doses (0.8 Gy in the BxPC-3 model), as shown in Table 2, and Figures S2-S4). Fig. 4B indicates the time required for tumor volumes to reach/not exceed the end-point of 400mm^3^.

**Figure 4.**
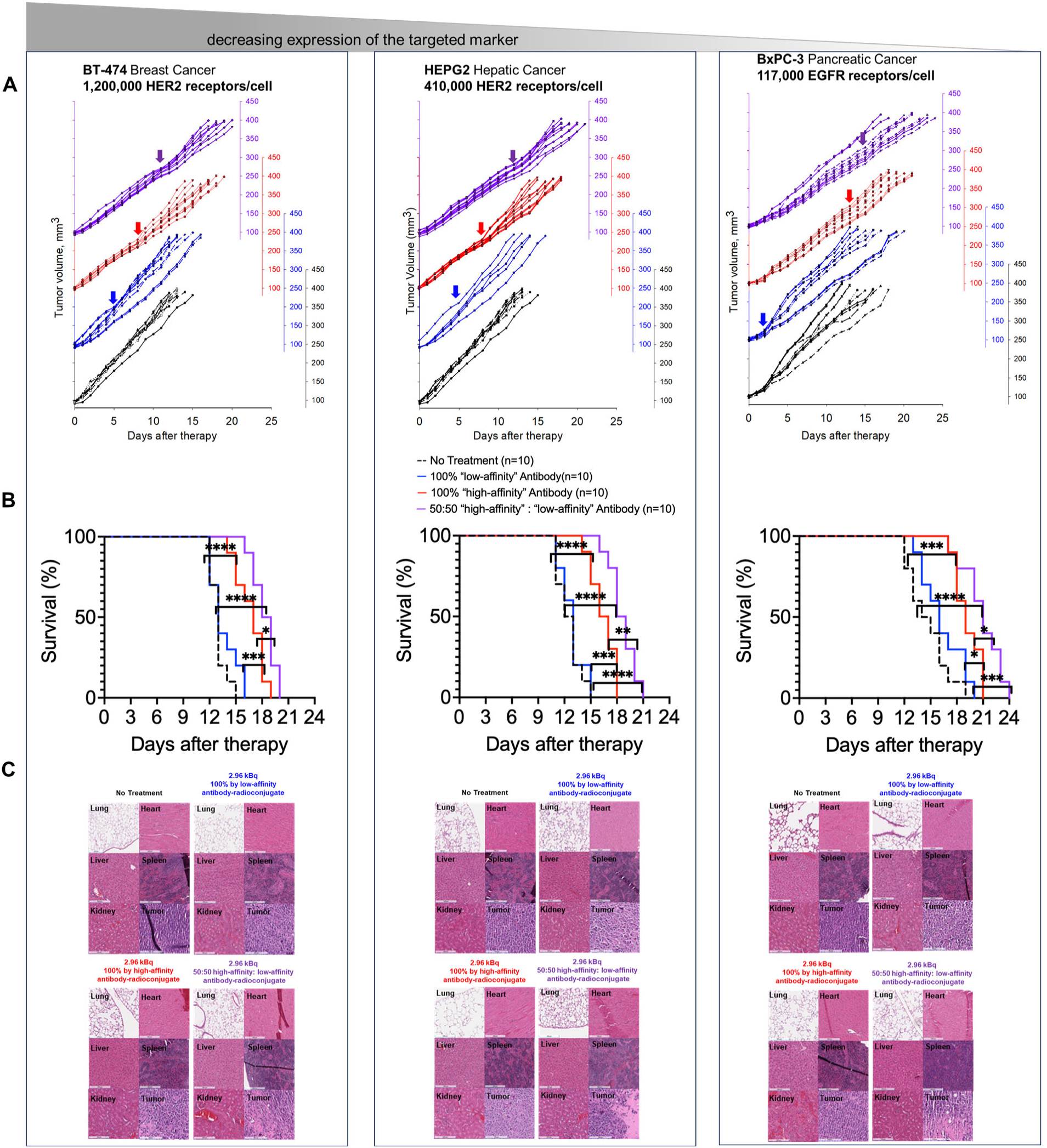
Independent of tumor origin, and the type and expression levels of the targeted markers on cancer cells, affinity cocktails of high-affinity and low-affinity actinium-225 antibody-radioconjugates (purple symbols, lines) were more effective in inhibiting tumor growth at the same total systemically injected activity compared to the high-affinity antibody-radioconjugate alone (red symbols/lines). (A) Growth inhibition of subcutaneous xenografts in NSG mice injected systemically with 2.96kBq actinium-225 delivered by each affinity form of antibody-radioconjugate alone and combined. Vertical arrows indicate the inflection points beyond which tumors exhibited increased growth rates. (B) Animal survival indicating the time required for tumor volumes to reach/not exceed the end-point of 400mm^3^. (C) H&E staining of tumors and critical off-target organs. Scale bar = 200µm.

In all groups, at the time of sacrifice, pathology assessment of excised tissues did not indicate noteworthy cardiac, renal or hepatic toxicities (Fig. 4C). Mild to moderate levels of extramedullary hematopoiesis (EMH) were observed in the liver and spleen across all three tumor models. The non-treated group exhibited the highest EMH levels in the liver, while EMH was reduced in treated conditions - with the lowest levels observed in the groups that received the affinity cocktails (animal weight measurements vs. time are plotted in Fig. S<u>8</u>).

## Discussion

Antibody-radioconjugates are being employed by most of the currently active clinical trials on TAT for the treatment of solid tumors both for tumors with high and moderate expression of the targeted markers (NCT04147819, (*13*)). In general, these antibody-radioconjugates have significant (high) binding affinity toward the targeted receptors (*1*). However, high affinity limits the antibodies’ penetration within solid tumors potentially resulting in heterogeneous α-particle tumor irradiation patterns, especially within those tumors with high expression levels of the targeted receptors, therefore compromising efficacy (*6,11*). In this proof-of-concept study that evaluated affinity cocktails of antibody-radioconjugates, we demonstrated that delivering a fraction of the same total administered activity by a less strongly binding form of (the same) antibody (targeting the same marker on tumor cells as the high-affinity antibody) results in better inhibition of tumor growth independent of tumor origin, and the type and expression levels of the targeted marker. Our measurements in spheroids, employed as surrogates of tumor avascular regions, supported the hypothesis that the *in vivo* observations of the better tumor growth inhibition by the affinity cocktails of antibody-radioconjugates, compared to the inhibition by the same total injected activity delivered only by the high-affinity antibody-conjugates, can be largely attributed to the better spreading of the delivered α-particle emitters within the solid tumor volume.

In this study, the TAT carriers employed were only antibody-radioconjugates of different affinity: this choice of carriers aimed to potentially facilitate clinical adaptation, since antibody-delivered TAT is already in the clinic and, this study shows that the approach of affinity cocktails is generalizable. Notably, on the mouse xenograft models studied, the extent of tumor growth inhibition did not generally align with the tumor absorbed doses, indicating the significance of augmenting the reported levels of tumor absorbed doses with the patterns of tumor irradiation by α-particles, as we previously demonstrated (*3,6,11*).

Regarding potential toxicities of the low(er)-affinity antibody-radioconjugates, we demonstrated that both the high-affinity and low-affinity trastuzumab-radioconjugates exhibited indistinguishable off-target uptake in mice (Fig. S2). This finding is possible to be also the case in humans and could play a critical role in addressing clinical translation considerations. We employed a simple, reproducible approach to decrease the antibody affinity (for low-affinity trastuzumab) by click chemistry selectively targeting the α-amino groups on the antibody paratopes, but, certainly, any other antibody-engineering approach can achieve the same result in tuning the antibodies’ affinity.

The different affinity antibody-radioconjugates were combined in the same injectate and were administered simultaneously. Asynchronous injection of each separate radioconjugate was not evaluated but could possibly increase both the tumor uptake and the penetration of the subsequently injected TAT form, since pre-irradiation of the tumor by the first radio-conjugate is possible to affect the viability and, thus, the effective “porosity” of solid tumors toward the following conjugate (*14*).

We employed an experimentally-informed first-principles diffusion-reaction mathematical model (a digital twin), that we trained on spheroids (*9,15*). The digital twin predicted the 70:30 activity split ratio between the high-affinity and low-affinity trastuzumab-radioconjugates as more fit than the 50:50 ratio, and the validity of this prediction was demonstrated in our studies (shown in Fig. 2). Briefly, the mathematical model (*15*), evaluated the activity split ratios between the two antibody-radioconjugates that maximize the number of cells within spheroids which, for the least possible incubating activity, receive lethal doses of actinium-225. The optimization is dependent on the spatiotemporal distributions of delivered α-particles by each antibody-radioconjugate (Fig. S5), and, critically, by the mean expression levels of the targeted marker by cancer cells comprising the spheroids.

The affinity cocktails of antibody-radioconjugates are conceptually different from previously reported cocktails of α-particle antibody-radioconjugates, where *each and every one of the antibodies in the cocktail are chosen to strongly bind to a (different) receptor/marker expressed on the surface of the same cancer cells* comprising the tumor (*16*). The idea behind the latter cocktails with both antibodies binding to a different marker, is to *collectively increase the activity delivered per cancer cell*, when none of the targeted markers is overexpressed by cancer cells. Our approach addresses the heterogeneous irradiation of solid tumors caused by high-affinity antibody-radioconjugates that exhibit limited tumor penetration.

In conclusion, this proof-of-concept study demonstrates the technical ease and the, potentially, straightforward clinical implementation of the affinity cocktails along with their general applicability to augment different types of antibody-radioconjugates, making them a platform technology for enhancing the efficacy of antibody-delivered TAT against solid tumors.

## Acknowledgements

The authors thank Dr. Denis Wirtz for sharing the BxPC-3 cell line. This study was partially supported by the MII/TEDCO and the Congressionally Directed Medical Research Programs, Award number HT9425-24-1-1003.

