## Supporting Information for "Guiding treatment response by spatiotemporal control of α-particle deposition in solid tumors: the case for ‘affinity cocktails’ of antibody-radioconjugates"

### Materials and Methods

#### *Materials*

Phosphate Buffered Saline (PBS), trisodium citrate dihydrate, anhydrous citric acid, poly(2-hydroxyethyl methacrylate) (polyHEMA) and 2-Amino-2-(hydroxymethyl)-1,3-propanediol (Trizma®) were purchased from Sigma-Aldrich (Atlanta, GA, USA). Trypsin and Matrigel™ were purchased from Corning (Corning, NY, USA), Ethylenediaminetetraacetic acid (EDTA) was purchased from Fisher Scientific (Pittsburgh, PA, USA), penicillin-streptomycin was from ThermoFisher Scientific (Waltham, MA, USA), S-2- (4-Isothiocyanatobenzyl)-diethylenetriamine pentaacetic acid (DTPA-SCN) and S-2-(4-isothiocyanatobenzyl)-1,4,7,10-tetraazacyclododecane-1,4,7,10-tetraacetic acid (DOTA-SCN) were from Macrocyclics (Dallas, TX, USA). Chelex® resin, chromatography and desalting columns from Bio-Rad (Hercules, CA, USA), syringe filters (0.22µm, Cat No. 76479-024) from VWR (Radnor, PA, USA), the Eagle's Minimal Essential Medium (EMEM) was purchased from Quality Biological (Gaithersburg, MD,USA) and, Hybricare™ and Roswell Park Memorial Institute (RPMI) medium were from ATCC (Manassas, VA, USA). The Fetal Bovine Serum (FBS) was from Omega Scientific (Tarzana, CA, USA), and the HER1-binding antibody Cetuximab was purchased from Eli Lilly (Indianapolis, IN, USA) while the HER-2 binding antibody Trastuzumab was from Genentech (San Francisco, CA, USA). The non-specific antibody Rituximab was purchased from the Johns Hopkins Pharmacy (Baltimore, MD, USA).

#### *Antibody radiolabeling and characterization*

For the radiolabeling, actinium-225 (or indium-111) dissolved in 0.2 N HCl was added to the chelator-conjugated antibody, suspended in 500µL of Tris-HCl buffer (or acetate buffer), and the reaction mixture was incubated at 37°C for one hour. Following this, the radiolabeled antibody was then purified using a 10DG column equilibrated with PBS at 1 mM, pH 7.4. Radiolabeling efficiency was calculated as the ratio of the measured radioactivity before and after the 10DG column. Radiochemical purity was evaluated using iTLC with 10 mM EDTA in water as the mobile phase [1]. Immunoreactivity of the (radio)labeled antibody was measured by incubating with cells on ice for 1 hour at 100:1 receptor:antibody ratio and, upon separation of non-cell-bound antibodies, by quantifying the radioactivity associated with cells relative to the total radioactivity added. The fraction bound was corrected for non-specific antibody binding to cells by evaluating, in parallel suspensions, the fraction of radiolabeled antibody bound to cells in the presence of 50x excess unlabeled antibody. The stability of the antibody radiolabeling was evaluated by adding the radiolabeled antibody to media at pH 7.4. Following 24 hours of incubation at 37°C, the antibody was passed through a 10DG column equilibrated with PBS at 1mM, pH 7.4, and the antibody fractions that were eluted from the column were collected. The stability of radiolabeling was then calculated as the ratio of the measured radioactivity associated with the antibody before incubation and after the completion of the 24-hour incubation.

*Clonogenic cell survival assay*

Cancer cells were plated in 6-well plates at 500,000 cells per well and were allowed to adhere overnight before being incubated with varying concentrations of activity (4.63, 9.25, 18.5, 37, 74 kBq/mL) delivered either in the form of [$^{225}$Ac]Ac-DOTA or by the indicated actinium-225 antibody-radioconjugates, at pH 7.4 for a duration of 6 hours. Post completion of incubation, the cells in each of the wells were washed with PBS at pH 7.4, gently scraped and resuspended at a concentration of 10,000 cells/mL in complete media at pH 7.4 and were then plated into tissue culture dishes at varying cell densities. Once cell colonies were observed (~10 doubling times), the media was removed, dishes were washed with water, and the colonies were fixed and stained using 6% (w/v) glutaraldehyde and 0.05% (w/v) Crystal violet, respectively, following which they were counted using a colony counter pen. The number of colonies counted for each of the treatment groups was then normalized by the number of colonies from the control group to obtain the survival fraction, while accounting for the plating efficiency [2].

*Spheroid formation, spatiotemporal distributions and treatment*

Spheroids were formed by seeding 3200 BT-474 and/or 1000 HEPG-2 cells per well in a PolyHEMA-coated 96-well round bottom plate, and centrifuging it for 10 minutes at 1023 rcf and 4° C. The formed spheroids were tracked for their size and were used upon reaching the desired size. The cell line BxPC3 did not form spheroids.

The spatiotemporal profiles of the specific and non-specific antibodies were evaluated by incubating 400 μm diameter spheroids with FITC-labeled specific Antibody (Trastuzumab) (0.06μM, ex/em: 494/518nm), with the FITC-labeled non-specific Antibody (Rituximab) (0.06μM, ex/em: 494/518nm) or with NHS-Fluorescein labeled low affinity (Trastuzumab) (0.06μM, ex/em: 494/518nm). Spheroids were harvested at various timepoints - during incubation with the carriers (uptake) and upon being transferred in fresh media (clearance) -, were flash frozen in cryochrome, mounted on OCT gel and sectioned at 20 μm thickness. The equatorial slices were then imaged using a confocal fluorescence microscope (Zeiss LSM 780 Confocal Microscope with filters, 10X objective, White Plains, NY). To generate the corresponding calibration curves, known concentrations of FITC-labeled antibody were measured, using the same microscope settings, in a quartz cuvette of 20 μm path-length. An in-house developed MATLAB erosion code was applied on the images of spheroid sections to calculate the average intensity/concentration within each 5 μm-wide concentric ring of the section which was then plotted vs. its radial position, and the time-integrated, radial concentrations were calculated using the trapezoidal rule.

In treatment studies, upon reaching a size of 400 μm in diameter, the spheroids were incubated with varying combinations of radiolabeled specific and non-specific antibody, for 24 hours, to roughly match their blood clearance kinetics in mice, at three different total radioactivity concentration of 0.5, 1.0, 3.0 kBq/mL. The total antibody mass for both the specific and non-specific antibodies, each, was maintained at 200 times excess of the HER2 receptors expressed by all cells comprising the spheroid. After incubation, the treated spheroids were transferred into fresh media (one spheroid per well in PolyHEMA-coated U-bottom plates), and the spheroid volume was monitored until an asymptote was reached for the volume of untreated spheroids. At that point, spheroids were transferred to adherent 96-well plates (one spheroid per well in cell culture-treated F-bottom plates), and, once the non-treated condition reached confluency, cells from each well were trypsinized and counted. The percent outgrowth/regrowth was evaluated as the number of cells counted in each treated condition normalized by the number of cells in the untreated condition.

**HEPG2: high-affinity Trastuzumab**

| | %IA/g | | | | | |
|---|---|---|---|---|---|---|
| Time (hours) | 1 | 8 | 24 | 48 | 72 | 96 |
| Heart | 5.60 ± 0.68 | 3.89 ± 0.27 | 1.86 ± 0.26 | 1.30 ± 0.14 | 1.13 ± 0.09 | 0.91 ± 0.05 |
| Lung | 0.03 ± 0.01 | 0.20 ± 0.03 | 0.20 ± 0.04 | 0.17 ± 0.02 | 0.17 ± 0.02 | 0.14 ± 0.01 |
| Liver | 7.00 ± 1.01 | 9.27 ± 0.83 | 12.40 ± 0.63 | 8.18 ± 0.15 | 4.00 ± 0.41 | 2.26 ± 0.15 |
| Spleen | 3.92 ± 0.42 | 7.65 ± 0.80 | 10.35 ± 0.38 | 8.72 ± 0.23 | 12.93 ± 0.41 | 6.77 ± 0.17 |
| Kidney | 5.65 ± 0.65 | 3.02 ± 0.42 | 1.88 ± 0.59 | 1.18 ± 0.13 | 1.33 ± 0.08 | 0.21 ± 0.03 |
| Intestine | 1.20 ± 0.09 | 0.92 ± 0.11 | 1.09 ± 0.12 | 0.89 ± 0.13 | 0.72 ± 0.05 | 0.42 ± 0.03 |
| Stomach | 1.09 ± 0.12 | 0.65 ± 0.06 | 0.37 ± 0.07 | 0.26 ± 0.05 | 0.20 ± 0.01 | 0.14 ± 0.01 |
| Bone | 0.09 ± 0.02 | 0.20 ± 0.03 | 0.21 ± 0.07 | 0.17 ± 0.02 | 0.07 ± 0.00 | 0.07 ± 0.02 |
| Muscle | 0.11 ± 0.02 | 0.11 ± 0.05 | 0.27 ± 0.06 | 0.12 ± 0.03 | 0.05 ± 0.00 | 0.15 ± 0.01 |
| Tumor | 1.97 ± 0.28 | 3.21 ± 0.54 | 3.75 ± 0.48 | 4.76 ± 0.53 | 2.83 ± 0.20 | 1.81 ± 0.11 |
| Blood | 15.64 ± 2.88 | 11.02 ± 0.81 | 9.34 ± 0.35 | 5.99 ± 0.13 | 4.05 ± 0.14 | 2.28 ± 0.01 |

**HEPG2: Low-affinity Trastuzumab**

| | %IA/g | | | | | |
|---|---|---|---|---|---|---|
| Time (hours) | 1 | 8 | 24 | 48 | 72 | 96 |
| Heart | 4.14 ± 0.24 | 2.65 ± 0.10 | 1.63 ± 0.14 | 1.20 ± 0.12 | 1.11 ± 0.05 | 1.02 ± 0.12 |
| Lung | 0.11 ± 0.07 | 0.13 ± 0.02 | 0.37 ± 0.15 | 0.18 ± 0.06 | 0.10 ± 0.04 | 0.19 ± 0.05 |
| Liver | 4.43 ± 0.37 | 7.54 ± 0.87 | 11.57 ± 0.66 | 7.30 ± 0.90 | 4.32 ± 0.53 | 3.38 ± 0.48 |
| Spleen | 2.26 ± 0.27 | 5.67 ± 0.51 | 9.27 ± 0.09 | 7.78 ± 1.33 | 9.73 ± 0.08 | 9.96 ± 1.68 |
| Kidney | 4.68 ± 0.05 | 3.14 ± 0.25 | 1.97 ± 0.27 | 1.18 ± 0.06 | 0.89 ± 0.18 | 0.27 ± 0.06 |
| Intestine | 3.55 ± 0.80 | 1.05 ± 0.20 | 0.60 ± 0.09 | 0.58 ± 0.05 | 0.22 ± 0.03 | 0.16 ± 0.04 |
| Stomach | 1.94 ± 0.08 | 0.27 ± 0.03 | 0.19 ± 0.02 | 0.05 ± 0.06 | 0.11 ± 0.09 | 0.10 ± 0.07 |
| Bone | 0.09 ± 0.21 | 0.07 ± 0.03 | 0.19 ± 0.07 | 0.10 ± 0.01 | 0.18 ± 0.01 | 0.10 ± 0.03 |
| Muscle | 0.14 ± 0.11 | 0.07 ± 0.04 | 0.10 ± 0.10 | 0.12 ± 0.05 | 0.09 ± 0.03 | 0.06 ± 0.03 |
| Tumor | 1.23 ± 0.15 | 1.86 ± 0.42 | 2.90 ± 0.41 | 3.43 ± 0.84 | 1.68 ± 0.54 | 1.06 ± 0.13 |
| Blood | 13.22 ± 2.61 | 9.78 ± 2.06 | 9.49 ± 1.73 | 5.05 ± 0.27 | 4.80 ± 1.11 | 2.92 ± 0.95 |

**HEPG2: model low-affinity antibody (Rituximab)**

| | %IA/g | | | | | |
|---|---|---|---|---|---|---|
| Time (hours) | 1 | 8 | 24 | 48 | 72 | 96 |
| Heart | 5.74 ± 0.81 | 3.96 ± 0.87 | 2.29 ± 0.72 | 1.44 ± 0.24 | 1.02 ± 0.19 | 0.83 ± 0.22 |
| Lung | 0.04 ± 0.01 | 0.05 ± 0.01 | 0.14 ± 0.01 | 0.20 ± 0.05 | 0.12 ± 0.03 | 0.08 ± 0.01 |
| Liver | 6.37 ± 0.91 | 8.29 ± 1.22 | 11.15 ± 0.73 | 9.54 ± 1.12 | 5.76 ± 0.65 | 4.33 ± 0.17 |
| Spleen | 4.77 ± 0.91 | 6.66 ± 0.95 | 8.32 ± 0.95 | 9.51 ± 0.37 | 10.00 ± 0.92 | 4.99 ± 0.29 |
| Kidney | 4.01 ± 0.95 | 6.15 ± 0.95 | 4.00 ± 0.52 | 1.96 ± 0.23 | 1.35 ± 0.22 | 0.63 ± 0.11 |
| Intestine | 0.92 ± 0.20 | 1.02 ± 0.14 | 1.15 ± 0.18 | 0.78 ± 0.15 | 0.41 ± 0.10 | 0.27 ± 0.03 |
| Stomach | 1.02 ± 0.12 | 0.67 ± 0.17 | 0.83 ± 0.23 | 0.49 ± 0.13 | 0.35 ± 0.03 | 0.23 ± 0.03 |
| Bone | 0.24 ± 0.05 | 0.35 ± 0.01 | 0.42 ± 0.08 | 0.31 ± 0.04 | 0.24 ± 0.03 | 0.14 ± 0.03 |
| Muscle | 0.25 ± 0.04 | 0.44 ± 0.08 | 0.60 ± 0.07 | 0.39 ± 0.06 | 0.28 ± 0.05 | 0.17 ± 0.04 |
| Tumor | 1.64 ± 0.35 | 2.69 ± 0.55 | 5.23 ± 0.33 | 3.12 ± 0.18 | 2.01 ± 0.23 | 0.96 ± 0.11 |
| Blood | 17.07 ± 2.02 | 13.95 ± 1.11 | 10.62 ± 0.66 | 6.66 ± 0.51 | 4.84 ± 0.42 | 2.64 ± 0.28 |

Table S1. Biodistributions of radiolabeled antibodies (as indicated on the tables) on the HEPG2 subcutaneous xenografts following systemic injection in mice.

**BT-474: (high-affinity) Trastuzumab**

| %IA/g | | | | | | |
|---|---|---|---|---|---|---|
| Time (hours) | 1 | 8 | 24 | 48 | 72 | 96 |
| Heart | 5.85 ± 1.01 | 4.13 ± 0.66 | 2.07 ± 0.32 | 1.36 ± 0.19 | 1.18 ± 0.17 | 0.99 ± 0.16 |
| Lung | 0.10 ± 0.03 | 0.25 ± 0.04 | 0.09 ± 0.05 | 0.19 ± 0.03 | 0.19 ± 0.03 | 0.19 ± 0.01 |
| Liver | 8.67 ± 1.91 | 10.16 ± 1.10 | 13.33 ± 0.71 | 10.35 ± 0.51 | 4.37 ± 0.31 | 2.97 ± 0.17 |
| Spleen | 4.55 ± 0.68 | 6.76 ± 0.96 | 8.43 ± 0.44 | 10.77 ± 0.30 | 11.61 ± 0.44 | 4.87 ± 0.26 |
| Kidney | 4.34 ± 0.83 | 6.57 ± 0.85 | 3.22 ± 0.57 | 1.24 ± 0.33 | 0.76 ± 0.08 | 0.63 ± 0.04 |
| Intestine | 1.52 ± 0.20 | 1.53 ± 0.32 | 0.97 ± 0.23 | 0.74 ± 0.13 | 0.48 ± 0.03 | 0.23 ± 0.02 |
| Stomach | 0.84 ± 0.07 | 0.57 ± 0.12 | 0.56 ± 0.08 | 0.37 ± 0.06 | 0.28 ± 0.02 | 0.15 ± 0.02 |
| Bone | 0.20 ± 0.03 | 0.10 ± 0.02 | 0.33 ± 0.08 | 0.20 ± 0.04 | 0.17 ± 0.02 | 0.12 ± 0.01 |
| Muscle | 0.19 ± 0.03 | 0.19 ± 0.06 | 0.30 ± 0.04 | 0.26 ± 0.03 | 0.19 ± 0.01 | 0.12 ± 0.01 |
| Tumor | 2.09 ± 0.50 | 3.43 ± 0.57 | 5.54 ± 0.42 | 9.99 ± 0.81 | 5.35 ± 0.45 | 2.68 ± 0.16 |
| Blood | 16.93 ± 2.44 | 11.77 ± 1.00 | 9.52 ± 0.42 | 6.36 ± 0.27 | 4.35 ± 0.21 | 2.65 ± 0.04 |

**BT-474: model low-affinity antibody (Rituximab)**

| %IA/g | | | | | | |
|---|---|---|---|---|---|---|
| Time (hours) | 1 | 8 | 24 | 48 | 72 | 96 |
| Heart | 6.20 ± 0.74 | 4.11 ± 0.75 | 2.47 ± 0.49 | 1.47 ± 0.21 | 1.02 ± 0.14 | 0.95 ± 0.10 |
| Lung | 0.10 ± 0.04 | 0.11 ± 0.03 | 0.15 ± 0.03 | 0.17 ± 0.02 | 0.15 ± 0.02 | 0.09 ± 0.02 |
| Liver | 6.26 ± 0.55 | 8.81 ± 0.91 | 12.75 ± 0.94 | 7.86 ± 0.49 | 4.22 ± 0.49 | 2.97 ± 0.24 |
| Spleen | 4.54 ± 0.59 | 6.57 ± 0.67 | 8.59 ± 0.85 | 9.59 ± 0.43 | 12.42 ± 0.52 | 5.60 ± 0.41 |
| Kidney | 4.76 ± 0.81 | 6.57 ± 0.74 | 3.69 ± 0.64 | 2.13 ± 0.52 | 1.54 ± 0.12 | 0.77 ± 0.12 |
| Intestine | 1.25 ± 0.15 | 1.81 ± 0.27 | 1.28 ± 0.16 | 0.91 ± 0.21 | 0.53 ± 0.10 | 0.33 ± 0.03 |
| Stomach | 0.87 ± 0.20 | 0.74 ± 0.18 | 0.92 ± 0.09 | 0.41 ± 0.12 | 0.45 ± 0.05 | 0.28 ± 0.04 |
| Bone | 0.46 ± 0.07 | 0.43 ± 0.08 | 0.53 ± 0.10 | 0.37 ± 0.04 | 0.24 ± 0.03 | 0.17 ± 0.03 |
| Muscle | 0.34 ± 0.10 | 0.46 ± 0.09 | 0.71 ± 0.17 | 0.47 ± 0.12 | 0.32 ± 0.05 | 0.19 ± 0.02 |
| Tumor | 1.84 ± 0.53 | 2.87 ± 0.35 | 4.53 ± 0.57 | 3.01 ± 0.39 | 1.65 ± 0.20 | 1.11 ± 0.11 |
| Blood | 14.63 ± 1.90 | 10.57 ± 1.01 | 8.32 ± 0.71 | 5.16 ± 0.60 | 3.35 ± 0.43 | 2.09 ± 0.26 |

Table S2. Biodistributions of radiolabeled antibodies (as indicated on the tables) on the BT-474 subcutaneous xenografts following systemic injection in mice.

**BxPC-3: (high-affinity) Cetuximab**

| %IA/g | | | | | | |
|---|---|---|---|---|---|---|
| Time (hours) | 1 | 8 | 24 | 48 | 72 | 96 |
| Heart | 6.35 ± 1.93 | 3.36 ± 1.03 | 1.66 ± 0.33 | 1.30 ± 0.16 | 1.10 ± 0.60 | 0.65 ± 0.13 |
| Lung | 0.09 ± 0.01 | 0.15 ± 0.03 | 0.18 ± 0.10 | 0.30 ± 0.05 | 0.26 ± 0.11 | 0.23 ± 0.09 |
| Liver | 7.47 ± 1.08 | 9.49 ± 1.24 | 12.30 ± 1.17 | 9.74 ± 2.18 | 4.70 ± 1.64 | 1.42 ± 0.49 |
| Spleen | 4.22 ± 0.73 | 8.91 ± 0.91 | 10.16 ± 1.87 | 13.60 ± 2.23 | 18.30 ± 2.58 | 9.03 ± 1.56 |
| Kidney | 4.56 ± 1.11 | 4.00 ± 0.40 | 2.54 ± 0.45 | 1.24 ± 0.37 | 1.32 ± 0.10 | 0.06 ± 0.02 |
| Intestine | 1.87 ± 0.73 | 3.38 ± 1.95 | 1.40 ± 0.13 | 1.28 ± 0.10 | 0.78 ± 0.20 | 0.16 ± 0.09 |
| Stomach | 1.41 ± 0.18 | 1.13 ± 0.41 | 0.67 ± 0.06 | 0.34 ± 0.04 | 0.29 ± 0.08 | 0.13 ± 0.09 |
| Bone | 0.15 ± 0.05 | 0.21 ± 0.13 | 0.11 ± 0.09 | 0.19 ± 0.09 | 0.04 ± 0.01 | 0.08 ± 0.01 |
| Muscle | 0.17 ± 0.10 | 0.22 ± 0.09 | 0.12 ± 0.08 | 0.18 ± 0.08 | 0.04 ± 0.01 | 0.09 ± 0.02 |
| Tumor | 1.75 ± 0.89 | 2.06 ± 1.55 | 2.79 ± 1.39 | 3.33 ± 1.72 | 1.67 ± 1.35 | 1.52 ± 1.10 |
| Blood | 14.70 ± 1.12 | 11.70 ± 0.65 | 9.11 ± 1.49 | 7.23 ± 0.13 | 4.46 ± 0.69 | 2.69 ± 2.00 |

**BxPC-3: model low-affinity antibody (Rituximab)**

| %IA/g | | | | | | |
|---|---|---|---|---|---|---|
| Time (hours) | 1 | 8 | 24 | 48 | 72 | 96 |
| Heart | 6.11 ± 1.02 | 4.46 ± 0.62 | 2.01 ± 0.36 | 1.55 ± 0.17 | 1.10 ± 0.18 | 0.96 ± 0.21 |
| Lung | 0.05 ± 0.01 | 0.20 ± 0.06 | 0.20 ± 0.04 | 0.19 ± 0.04 | 0.21 ± 0.03 | 0.14 ± 0.02 |
| Liver | 7.18 ± 1.73 | 9.60 ± 1.70 | 12.40 ± 0.66 | 9.43 ± 1.11 | 5.19 ± 0.61 | 3.26 ± 0.17 |
| Spleen | 5.59 ± 1.04 | 7.05 ± 1.11 | 9.36 ± 0.47 | 10.35 ± 0.31 | 9.10 ± 0.51 | 5.52 ± 0.71 |
| Kidney | 3.66 ± 0.53 | 5.17 ± 0.90 | 4.10 ± 0.72 | 1.82 ± 0.39 | 0.90 ± 0.11 | 0.60 ± 0.08 |
| Intestine | 0.78 ± 0.23 | 1.17 ± 0.33 | 1.33 ± 0.20 | 0.61 ± 0.05 | 0.32 ± 0.04 | 0.15 ± 0.03 |
| Stomach | 0.92 ± 0.11 | 0.65 ± 0.14 | 0.63 ± 0.05 | 0.40 ± 0.06 | 0.36 ± 0.03 | 0.17 ± 0.01 |
| Bone | 0.22 ± 0.03 | 0.25 ± 0.06 | 0.36 ± 0.05 | 0.25 ± 0.03 | 0.16 ± 0.02 | 0.10 ± 0.02 |
| Muscle | 0.23 ± 0.03 | 0.24 ± 0.06 | 0.35 ± 0.02 | 0.30 ± 0.04 | 0.22 ± 0.02 | 0.16 ± 0.02 |
| Tumor | 1.86 ± 0.21 | 2.38 ± 0.17 | 4.41 ± 0.35 | 2.45 ± 0.23 | 1.68 ± 0.10 | 1.17 ± 0.12 |
| Blood | 15.73 ± 2.44 | 9.82 ± 1.13 | 7.65 ± 0.52 | 5.48 ± 0.35 | 2.98 ± 0.27 | 2.02 ± 0.11 |

Table S3. Biodistributions of radiolabeled antibodies (as indicated on the tables) on the BxPC-3 subcutaneous xenografts following systemic injection in mice.

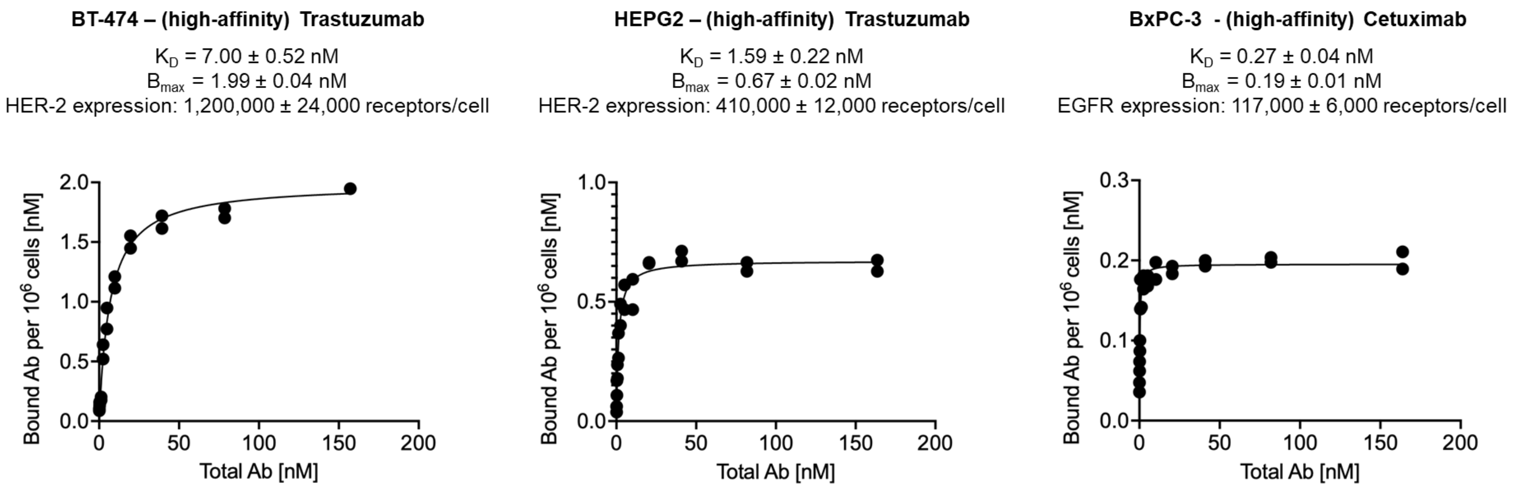


Figure S1. Binding isotherms of the (high-affinity) trastuzumab and cetuximab on the corresponding cancer cells lines employed in this study

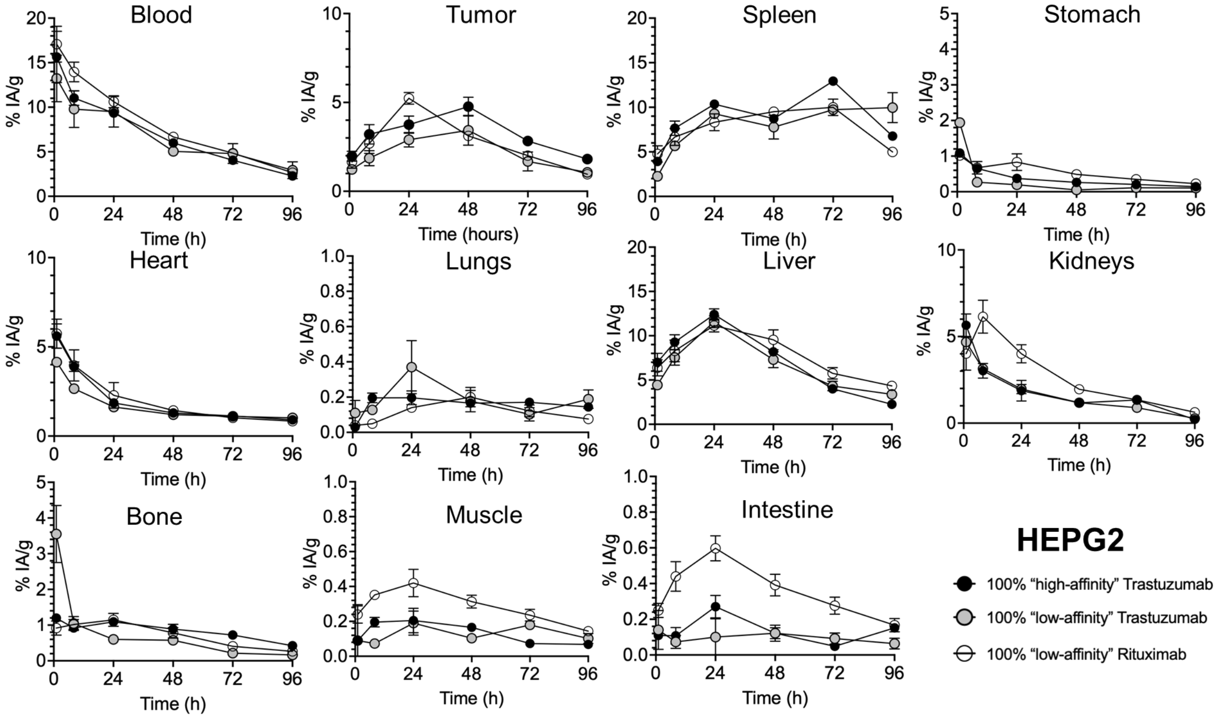


Figure S2. Biodistributions of radiolabeled antibodies (from table S1) on the HEPG2 subcutaneous xenografts following systemic injection in mice.

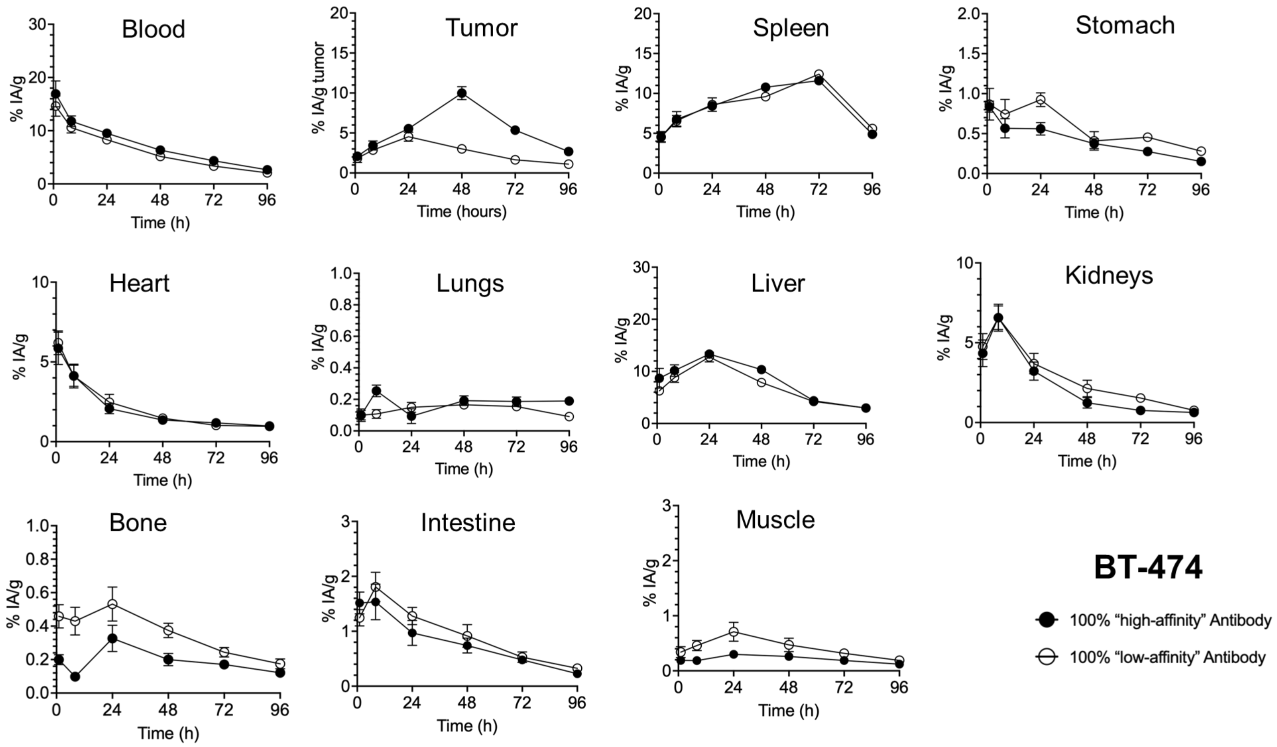


Figure S3. Biodistributions of radiolabeled antibodies (from table S2) on the BT-474 subcutaneous xenografts following systemic injection in mice. High-affinity antibody: (high-affinity) Trastuzumab; low-affinity antibody: model low/no affinity Rituximab.

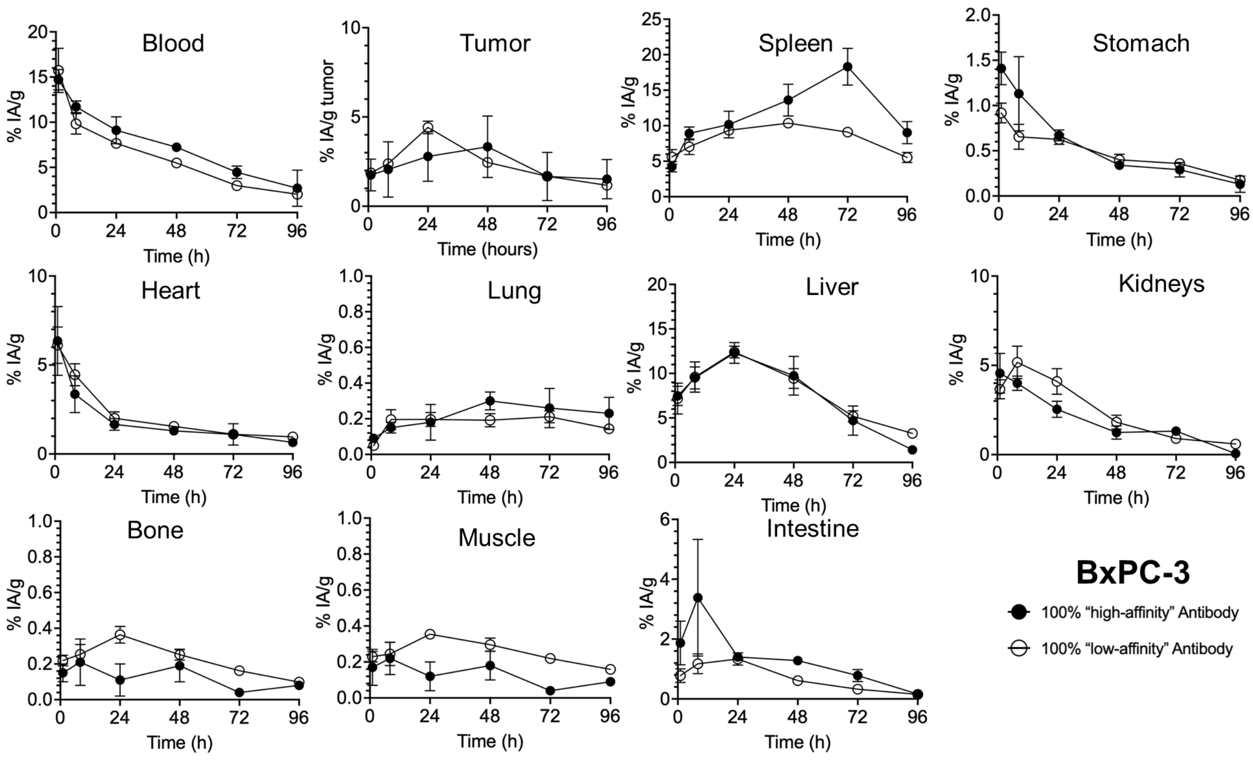


Figure S4. Biodistributions of radiolabeled antibodies (from table S3) on the Bx-PC3 subcutaneous xenografts following systemic injection in mice. High-affinity antibody: (high-affinity) Cetuximab; low-affinity antibody: model low/no affinity Rituximab.

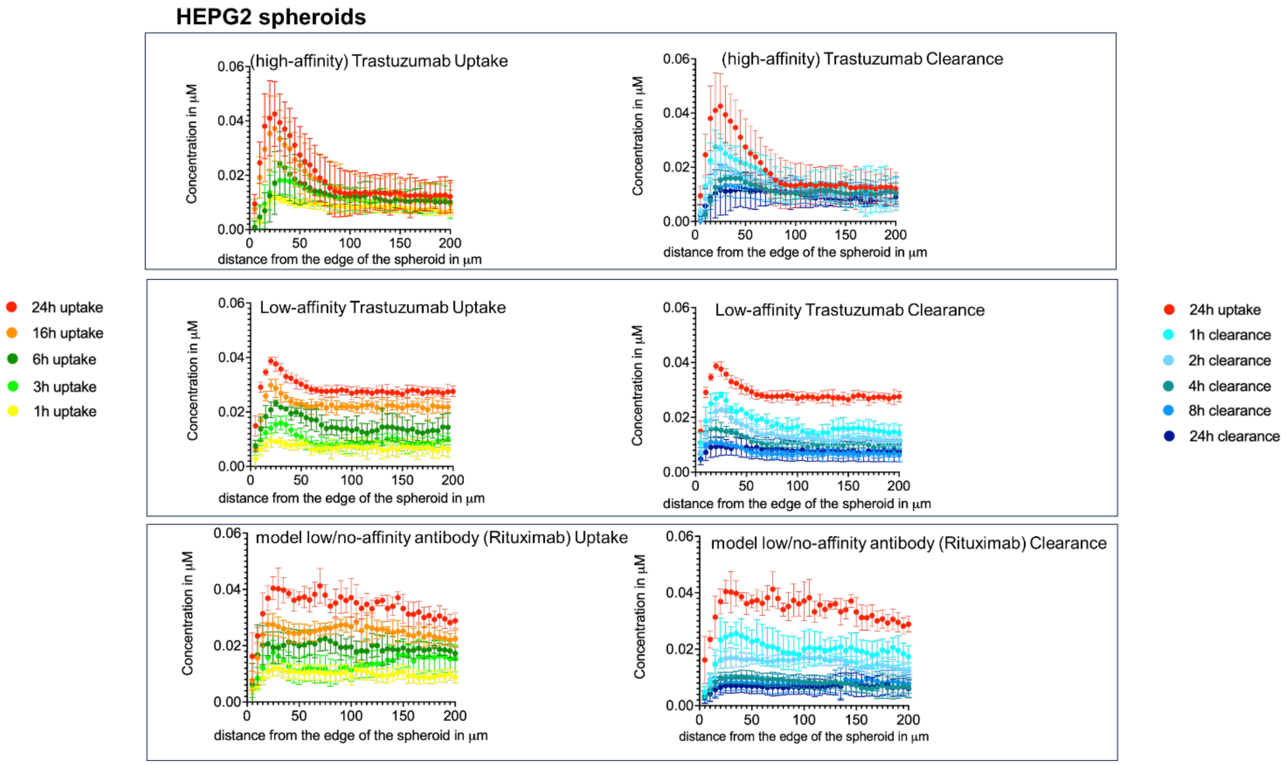


Figure S5. Spatiotemporal profiles of fluorescently-labeled antibodies (as indicated) on HEPG2 spheroids.

Individual Mouse Weights-HEPG2 Study with high and low affinity Trastuzumab

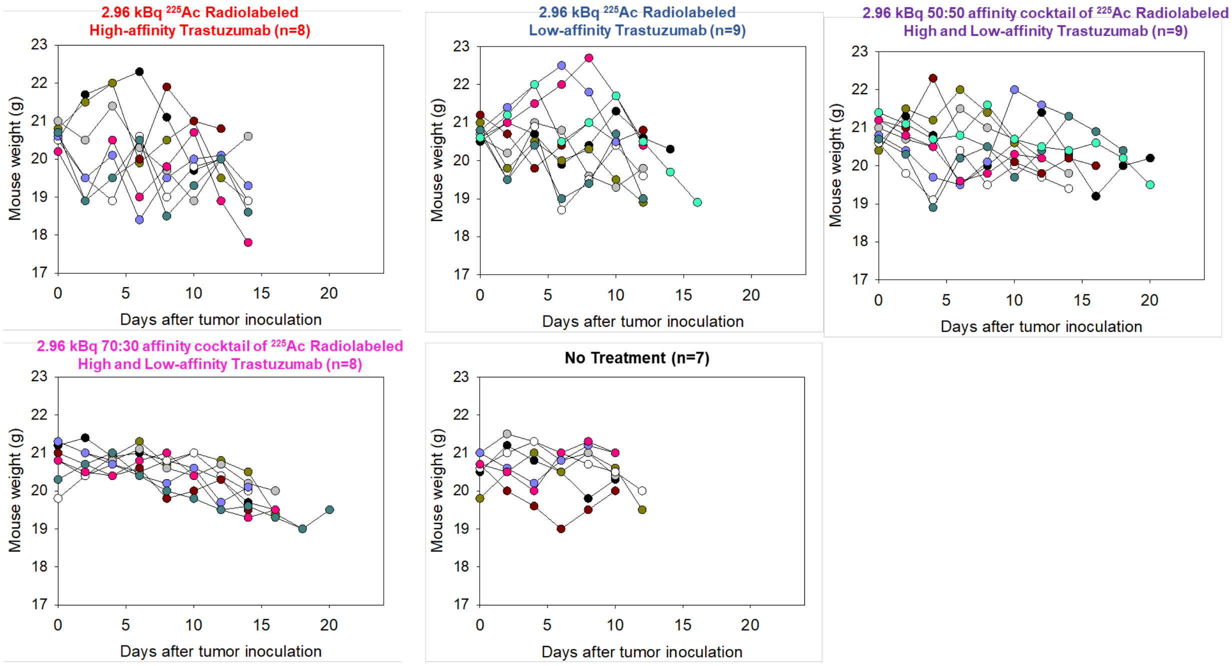


Figure S6. Mouse weights of the study shown on Figure 2.

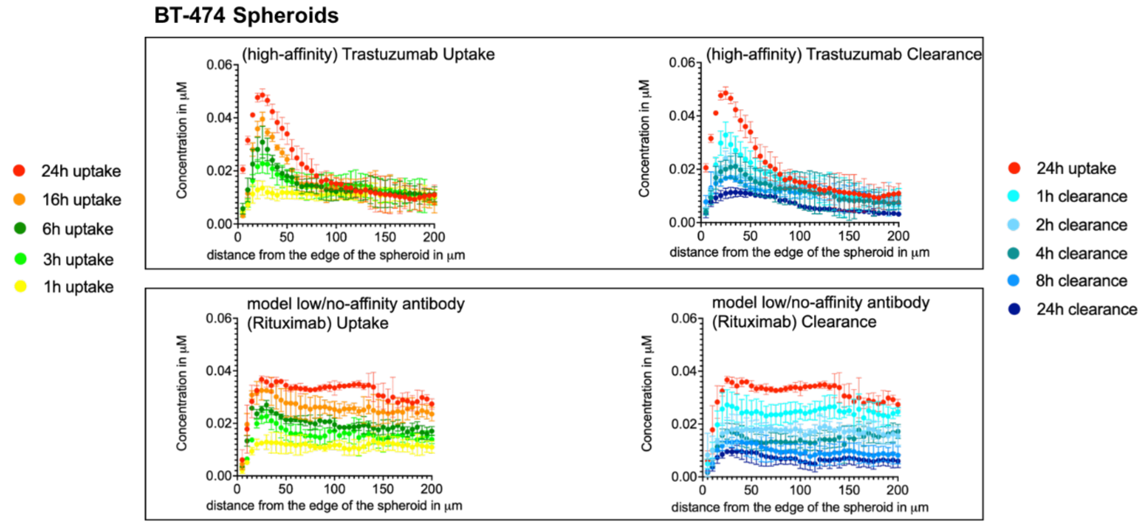


Figure S7. Spatiotemporal profiles of fluorescently-labeled antibodies (as indicated) on BT-474 spheroids.

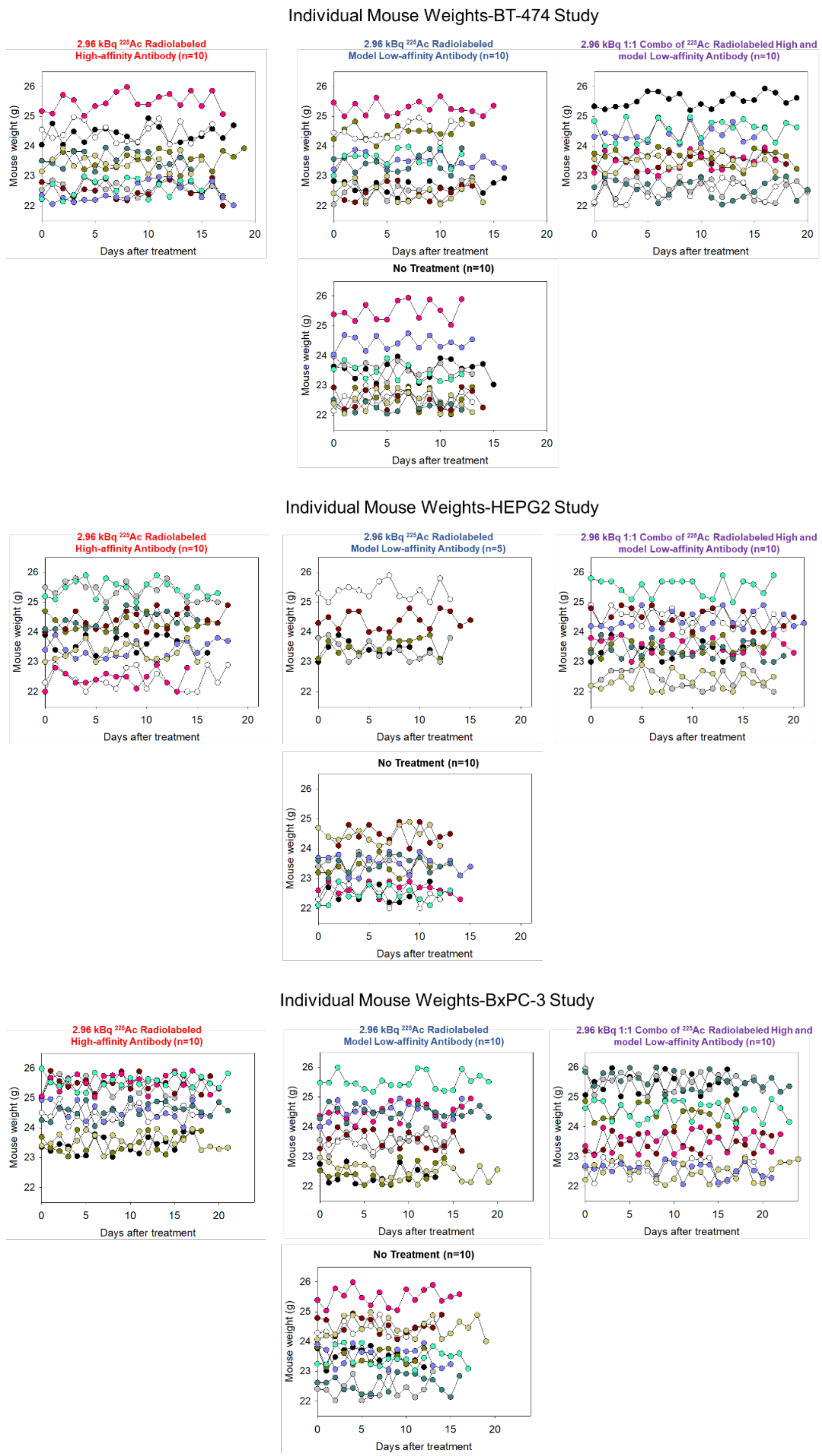


Figure S8. Mouse weights of the study shown on Figure 4.

*References*


1. McDevitt, M.R., et al., *Design and synthesis of actinium-225 radioimmunopharmaceuticals.* Appl Radiat Isot, 2002. **57**: p. 841-847.
2. Franken, N.A.P., et al., *Clonogenic assay of cells in vitro.* Nature Protocols, 2006. **1**(5): p. 2315-2319.
